# A *de novo* FtsA binder blocks late divisome recruitment, unmasking a hidden constriction mechanism

**DOI:** 10.64898/2026.11.15.766819

**Authors:** Jaana Männik, Jai M. Mehta, Jerry M. Parks, Jaan Männik

**Author notes:** **Corresponding author:** Jaan Männik. **Notice. Not for publication:** This manuscript has been authored by UT-Battelle, LLC under Contract No. DE-AC05-00OR22725 with the U.S. Department of Energy. The United States Government retains and the publisher, by accepting the article for publication, acknowledges that the United States Government retains a non-exclusive, paid-up, irrevocable, world-wide license to publish or reproduce the published form of this manuscript, or allow others to do so, for United States Government purposes. DOE will provide public access to these results of federally sponsored research in accordance with the DOE Public Access Plan (http://energy.gov/downloads/doe-public-access-plan).

## Abstract

Bacterial cytokinesis requires assembly of a multiprotein divisome complex, but the molecular event triggering its midcell recruitment and activity remains unclear. Recent work suggests a transition in the oligomeric structure of the conserved actin homolog FtsA may activate the divisome complex in *Escherichia coli*. Here, we test this model using a *de novo* designed protein binder, L125, predicted to block longitudinal FtsA oligomerization. L125 localizes to the Z-ring and inhibits cell division by blocking recruitment of late divisome proteins while enhancing recruitment of early division proteins. These findings implicate the FtsA longitudinal oligomerization interface in late divisome recruitment. Unexpectedly, L125-treated cells form broad, shallow constrictions that fail to complete division. These constrictions occur despite loss of detectable late-divisome localization and are consistent with ZipA-dependent preseptal cell-wall synthesis, previously not known to cause constrictions. This alternative constriction pathway may reinforce the future division site before activation of the canonical division machinery.

## Introduction

A central unresolved question in bacterial cytokinesis is how this process is regulated during the cell cycle ^1–3^. Cell division in most bacteria begins with the assembly of the Z-ring. The Z-ring is organized around FtsZ protofilaments that are tethered to the cytoplasmic membrane by membrane-associated linker proteins. In *Escherichia coli*, the principal linkers are FtsA and ZipA, both of which are essential in wild-type cells ^2,4,5^, although why two different linkers are necessary remains unclear.

Despite being the first step in bacterial cytokinesis, the formation of the Z-ring in *E. coli* does not immediately trigger constriction, which begins approximately one-quarter of the cell cycle later under several growth conditions ^6,7^. Before or at the time of the onset of constriction, so-called late-divisome proteins, the divisome, are recruited. The molecular events that occur during this delay and determine the recruitment of the late divisome proteins remain only partially understood.

Before constriction starts, FtsEX and the essential late divisome proteins FtsK, FtsQLB, FtsWI, and FtsN are recruited in largely hierarchical order to the Z-ring ^8,9^. FtsWI forms the core septal peptidoglycan (PG) synthase, which associates with FtsQLB to form the FtsWIQBL core divisome complex ^10,11^, whereas FtsN is the last essential division protein to be recruited ^12–14^. Although FtsN arrives near the time of constriction and has often been considered a trigger or activator of septal synthesis ^12,15–18^, its abundance is not rate-limiting for constriction ^7,19^. By contrast, FtsZ abundance can become rate-limiting at higher growth rates, suggesting that accumulation of FtsZ to a threshold level is needed to trigger the constriction formation ^19–22^.

Because FtsZ is cytoplasmic, the signal linking its accumulation at the Z-ring to divisome activation must be transmitted through membrane-associated divisome proteins. FtsA is likely to mediate this coupling because it both tethers FtsZ to the membrane and interacts with late divisome proteins ^5,23–26^. FtsA can adopt multiple oligomeric states, including 12-mer minirings, arcs ^27^, and actin-like antiparallel filaments ^28,29^. The finding that FtsA mutants with reduced or altered self-interaction can bypass ZipA and divide at shorter cell lengths led to an original model in which reduced FtsA oligomerization promotes late-divisome recruitment ^30^, potentially through the conversion of 12-mer FtsA minirings into shorter arcs or monomers by FtsZ protofilaments ^1,27^. By contrast, recent studies show that the active divisome is associated with antiparallel FtsA filaments ^28,29^.

Here, we address how FtsA interacts with and recruits the late divisome complex at the onset of constriction. To that end, we designed a *de novo* protein binder, L125, using the BindCraft pipeline ^31^ to target the longitudinal FtsA interface and interfere with FtsA-FtsA association. L125 inhibits cell division and localizes to the Z-ring, consistent with its *in vivo* binding to FtsA. It blocks recruitment of late divisome proteins, beginning with FtsK, while enhancing midcell localization of early divisome proteins, suggesting that the FtsA longitudinal interface is required for the recruitment of late divisome proteins. Unexpectedly, L125-expressing cells still form broad, shallow constrictions that develop slowly and fail to complete division. These constrictions form without detectable involvement of the core divisome complex. Instead, the data are consistent with ZipA-dependent preseptal PG synthesis, a pathway previously implicated in cell wall synthesis before invagination ^32–34^.

## Results

### Design of *de novo* protein binders targeting FtsA-FtsA interfaces

To perturb FtsA oligomerization, we designed binders that target either the lateral interface between antiparallel protofilaments or the longitudinal interface between successive FtsA monomers. Because the available *E. coli* FtsA structure contained only a single filament, we generated an *E. coli* FtsA filament model based on the antiparallel double-filament structure of *Vibrio maritimus* FtsA reported by Nierhaus *et al.* ^28^. The resulting lateral FtsA-FtsA interface was consistent with experimental observations by Perkins *et al.* ^29^. We then performed three hotspot-targeted BindCraft runs using sets of nine, five, and three exposed hydrophobic residues (see Methods; Supplementary Data 1, Hotspot-targeted). The nine-residue set targeted both interfaces, while the five- and three-residue sets targeted the lateral and longitudinal interfaces, respectively. Although several designs progressed to the fourth stage of the BindCraft pipeline and showed favorable design metrics, none passed all default computational filters. We nevertheless selected three of the highest-ranking candidates for experimental testing (Supplementary Table 1), but none altered cell length or showed detectable midcell localization when expressed as mNG fusions.

We therefore broadened the design strategy by removing predefined hotspot residues. An additional BindCraft run yielded 18 designs (Supplementary Data 1, Hotspot-free) that passed all default computational filters and converged on neighboring binding sites contacting both the IA and IIB domains of FtsA (Supplementary Fig. 1a). One design, L125, showed a broader interaction footprint across IIB and was predicted to block longitudinal contacts with an adjacent FtsA subunit (Fig. 1a,b; Supplementary Fig. 1a,b). L125 and three additional filter-passing longitudinal interface designs were tested (Supplementary Table 1). Only L125 altered cell length and localized to midcell.

**Fig. 1.**
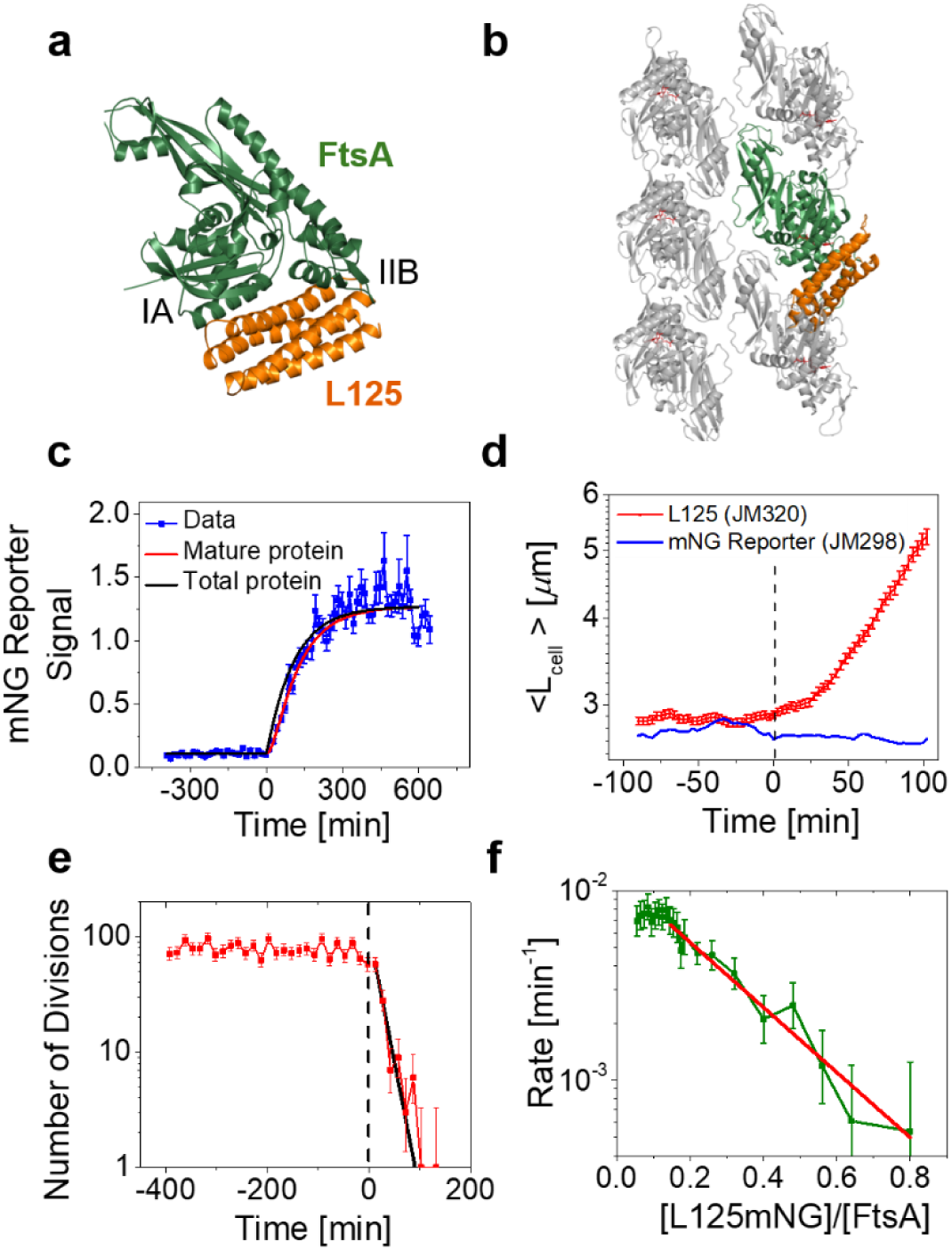
A *de novo*-designed protein binder, L125, targets the FtsA longitudinal interface and inhibits cell division in *E. coli*. **a** Structural model of L125 (orange) bound to FtsA (green) based on BindCraft ^31^. **b** The FtsA-L125 binding model mapped onto an antiparallel FtsA double-filament model (grey) based on PDB 7Q6F ^28^. **c** mNG reporter fluorescence (strain JM298) used as a proxy for L125 expression during protein induction. The red line is a fit to the mature fluorescent mNG concentration, and the black line is the predicted total mNG concentration after correction for fluorophore maturation (see Methods, *Determination of L125 concentration*). The concentration is in units of FtsA concentration in the cell population. **d** Population-average cell length, < *L_cell_* >, as a function of time after induction of expression of the L125 binder (red, strain JM320) and mNG reporter (blue, strain JM298). **e** Number of cell divisions per 15-min interval for the L125-expressing strain as a function of time. The black line is a fit to an exponential with a characteristic time of 19 min. **f** The population division rate as a function of [L125-mNG]/[FtsA] (strain JM344). Red line is an exponential fit for [*L*125*mNG*]/[*FtsA*] > 0.15, with a characteristic decay parameter 0.25. Error bars are s.e.m. for all panels except for e and f, where they are 68% Poisson confidence intervals. The number of cells analyzed is *N* = 4239 for JM298, *N* = 1596 for JM320 and *N* = 2127 for JM344.

### L125 halts cell division at substoichiometric levels relative to FtsA

L125 consists of four α-helices predicted to interact with helix 1 of the FtsA IA domain and additionally contact domain IIB through several non-covalent interactions (Supplementary Fig. 1b,c). When induced from a low-copy-number plasmid to approximately 140% of the native FtsA level, L125 caused rapid filamentation relative to a control strain expressing mNG from the same plasmid backbone (Fig. 1c,d; Supplementary Fig. 2a-c). Even without induction, when expressed at approximately 10% of the FtsA level, L125 increased average cell length by about 5% (Fig. 1d; Supplementary Fig. 2b). Given an estimated cellular FtsA concentration of approximately 1 µM under these conditions ^35^, about 100 nM L125 was sufficient to perturb division measurably.

The number of division events per 15-min interval remained approximately constant during the 400 min before induction but decreased exponentially afterward, with a characteristic time of about 20 min; no divisions were detected after 100 min (Fig. 1e). During this interval, reporter-based calibration indicated that the average L125 concentration increased from approximately 10% to 80% of the FtsA level (Fig. 1c), suggesting that a bulk [L125]/[FtsA] ratio below 1 is sufficient to stop division.

To quantify cell-to-cell variability in expression, we fused mNG to the C-terminus of L125 and expressed the fusion from the same plasmid backbone. L125-mNG inhibited division similarly to unlabeled L125 (Supplementary Fig. 2d), indicating that the tag did not substantially alter activity. L125-mNG fluorescence enabled the determination of its concentration in individual cells at division. The population-average concentration initially increased similarly to that of the mNG reporter, but later decreased rather than reaching the reporter plateau because shorter, lower-expressing cells were preferentially retained in mother-machine channels (Supplementary Fig. 2e). The fluorescence fusion nevertheless allowed L125-mNG concentration to be assigned to each cell at the moment of division, avoiding this population-sampling bias in the rate-versus-concentration analysis.

The population-average division rate was nearly constant for [L125-mNG]/[FtsA] < 0.15 and then decreased approximately exponentially with a characteristic concentration scale of 0.25 (Fig. 1f). Thus, a relatively small increase in binder abundance above this threshold produced a large reduction in division rate. No divisions were observed for [L125-mNG]/[FtsA] > 0.8. Thus, L125 and L125-mNG can completely inhibit division at substoichiometric levels relative to FtsA.

### L125-mNG exhibits FtsA-like Z-ring localization, consistent with its binding to FtsA

To investigate whether L125 inhibits division by binding to FtsA, we compared its localization with that of FtsA, measured independently using FtsA-GFP ^36^. At its basal expression level, corresponding to approximately 10% of the native FtsA abundance, L125-mNG caused only a minor division defect and localized predominantly to midcell throughout the cell cycle (Fig. 2a). Its cell-cycle-dependent midcell localization resembled that of FtsA-GFP, although its peak midcell excess fraction was lower (34% versus 53%; Fig. 2b). For brevity, we hereafter refer to the excess fraction and excess amount at midcell as the midcell fraction and midcell amount, respectively. These quantities estimate the fraction and amount of molecules associated with the Z-ring. In contrast, the localization pattern of L125-mNG differed from that of Ypet-FtsN, which accumulated at midcell only late in the cell cycle, and from that of the Z-ring-associated protein ZapA-mCherry (Supplementary Fig. 3a,b). Together, these observations are consistent with L125-mNG localization being determined by binding to FtsA.

**Fig. 2:**
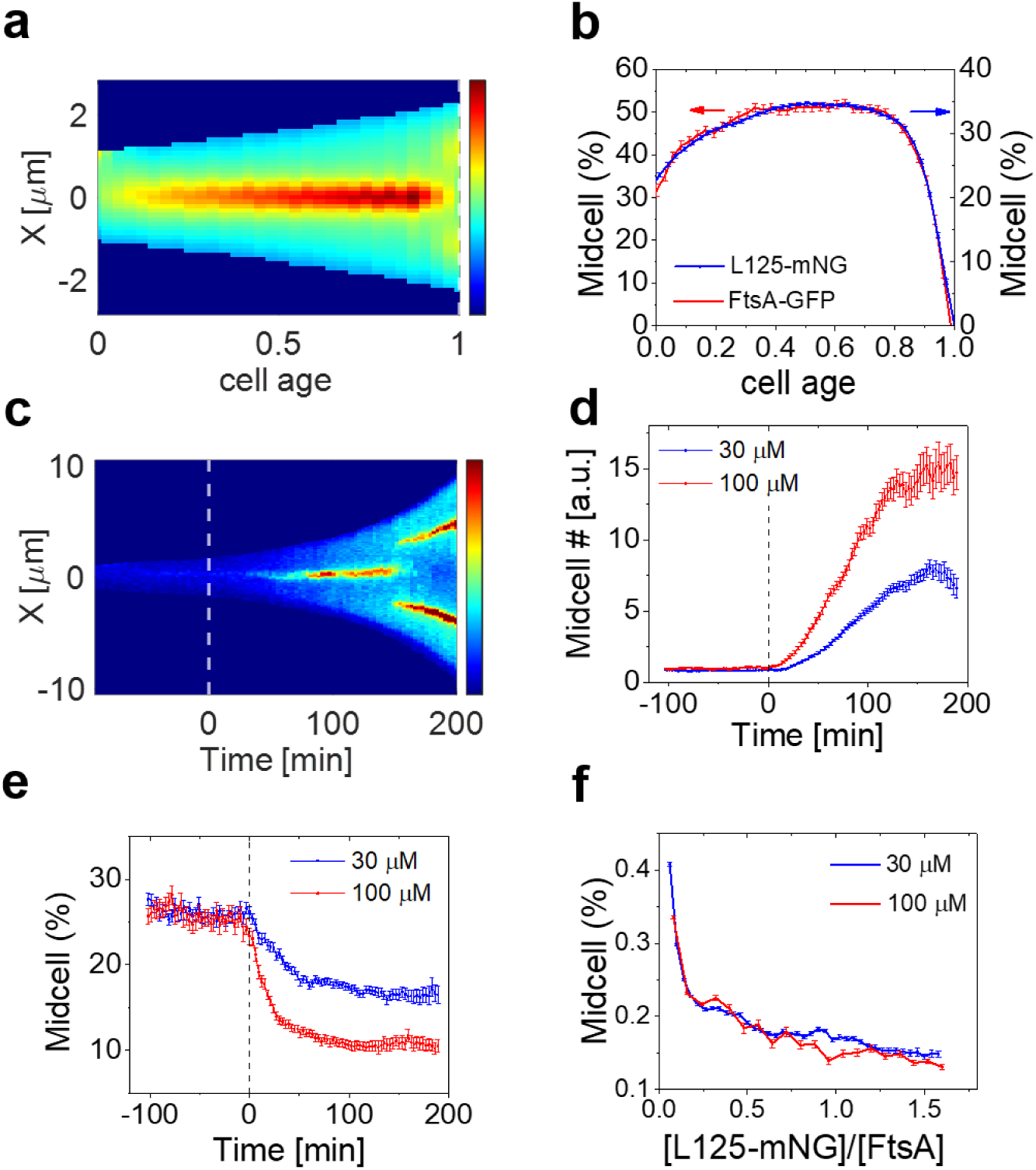
L125-mNG localizes to the Z-ring, similar to FtsA, and causes only minor perturbation to the Z-ring. **a** Kymograph of L125-mNG for the population of cells (*N*=515, strain JM344). L125-mNG is expressed at its basal level (not induced). The horizontal axis is the normalized cell age from birth (0) to the next division (1). The vertical axis is the distance, X, from the cell middle. **b** L125-mNG (right axis, *N*=515) and FtsA-GFP expressed from an additional gene copy (left axis, *N*=494, strain JM181) fractions in the Z-ring as a function of cell age. **c** Representative kymograph of a single cell expressing L125-mNG. Time is shown relative to L125-mNG induction by IPTG addition at 0 min, indicated by the white dashed line. **d** L125-mNG numbers at midcell (in the Z-ring) as a function of time after induction with 30 µM (blue) or 100 µM IPTG (red). **e** Fraction of L125-mNG at midcell as a function of time after induction. **f** Fraction of L125-mNG at midcell as a function of L125-mNG concentration expressed in units of FtsA concentration in the cell. *N* = 2127 for measurement with 30 μM and *N* = 827 with 100 μM induction. Error bars in all panels are s. e. m.

After induction, L125-mNG initially remained localized at midcell. However, as cells elongated to approximately 10 µm, the midcell band disappeared, and two bands emerged near the quarter positions (Fig. 2c; Supplementary Fig. 3c), similar to the splitting of early Z-ring protein bands reported in other division-defective *E. coli* strains ^37,38^. The midcell amount of L125-mNG increased after induction, with a larger increase at the higher inducer concentration (Fig. 2d), whereas the fraction of total cellular L125-mNG at midcell decreased (Fig. 2e,f). Even before band splitting, the decreasing midcell fraction is consistent with progressive saturation of a limited pool of L125-mNG binding sites at midcell: as the cellular concentration of L125-mNG increases, its increasing fraction remains cytosolic rather than associating with the Z-ring.

### L125 blocks the recruitment of late divisome proteins but enhances the accumulation of early divisome components

To investigate how FtsA oligomerization affects the recruitment of the other divisome proteins, we examined fluorescent fusions to selected divisome proteins. The early proteins FtsX and ZipA and the Z-ring-associated protein ZapA remained localized at midcell in the presence of L125 (Fig. 3a). In contrast, the essential late divisome proteins FtsK, FtsB, FtsI, and FtsN were not recruited (Fig. 3b). The remaining essential late divisome proteins FtsQ, FtsL, and FtsW, which were not tested, are part of the FtsBQL and FtsWI complexes. Because FtsB and FtsI were not recruited, FtsQ, FtsL, and FtsW also should not have been recruited. Thus, L125 preserved recruitment of the tested early divisome components, including FtsEX, while blocking recruitment of the late essential divisome proteins.

**Fig. 3.**
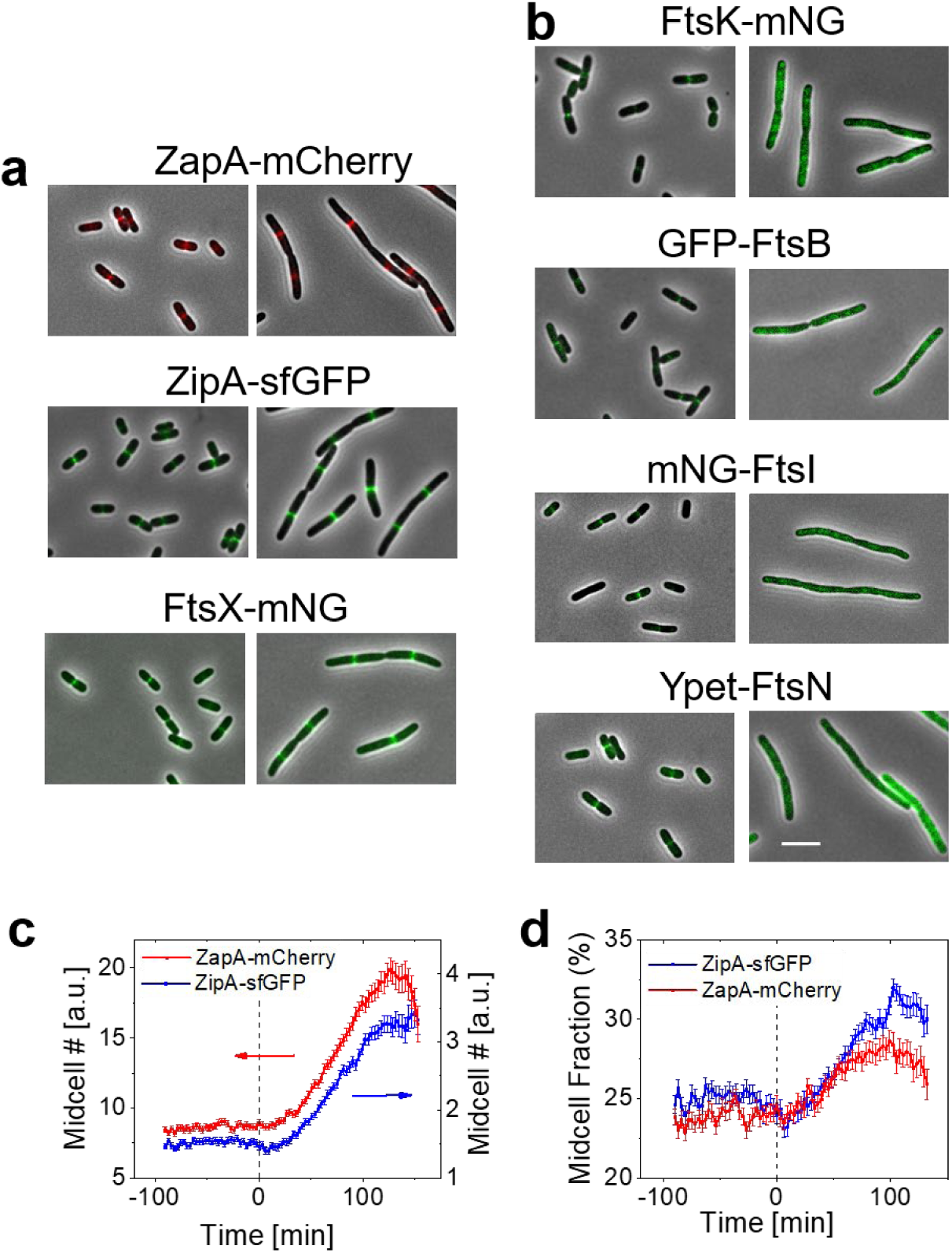
L125 does not block recruitment of early divisome proteins but blocks localization of essential late divisome proteins. **a** Representative phase-contrast and fluorescence image overlays showing localization patterns of fluorescent fusions to early divisome proteins before L125 induction (left panels) and 3 h after induction with 30 μM IPTG (right panels). **b** As in a, for a set of essential late divisome proteins. Scale bar, 5 μm (shown in b; applies to a,b). **c** Midcell numbers of ZipA-sfGFP (blue, left axis) and ZapA-mCherry (red, right axis) as a function of time after L125 induction. **d** Midcell fraction of ZipA-sfGFP and ZapA-mCherry signal after L125 induction.

Because early Z-ring components remained at midcell, we examined how their accumulation changed after L125 induction at two different IPTG concentrations, 30 and 100 μM. The midcell amounts of ZipA-sfGFP and ZapA-mCherry initially increased (Fig. 3c). Their relative midcell fractions also increased initially before their band split into two ¼ position bands (Fig. 3d). L125 thus enhances the accumulation of these two early divisome proteins. This finding contrasts with the decreasing midcell fraction of L125-mNG (cf. Fig. 2e). Thus, the reduction in L125-mNG midcell fraction cannot be explained by loss of early divisome components from the Z-ring. Instead, the finding further supports the notion that L125-mNG progressively saturates its FtsA binding sites as its concentration rises.

### Formation of broad constriction zones in the absence of late divisome localization

Although filamentous L125-expressing cells failed to complete division following the loss of late divisome recruitment, they nevertheless developed broad, shallow constrictions (Fig. 4a, Movie M1). These constrictions were not remnants of division events aborted by L125-mNG, because they began to appear only approximately 50–75 min after binder induction (Fig. 4b,c). The increase in the midcell phase-contrast signal coincided with a decrease in cell diameter at midcell (Fig. 4c), determined by fitting the fluorescence intensity profile across the short axis of the cell (Supplementary Fig. 4a). Broad constrictions also appeared at the quarter positions after the L125-mNG band split into two (Fig. 4d; Supplementary Fig. 4b–d). These secondary constrictions were weaker, consistent with the relatively short lifetime of the quarter-position L125-mNG bands, which subsequently reorganized into additional bands. Thus, constrictions can still form when L125-mNG blocks cell division, but they progress slowly and fail to complete before new division sites emerge at the cell quarter positions.

**Fig. 4.**
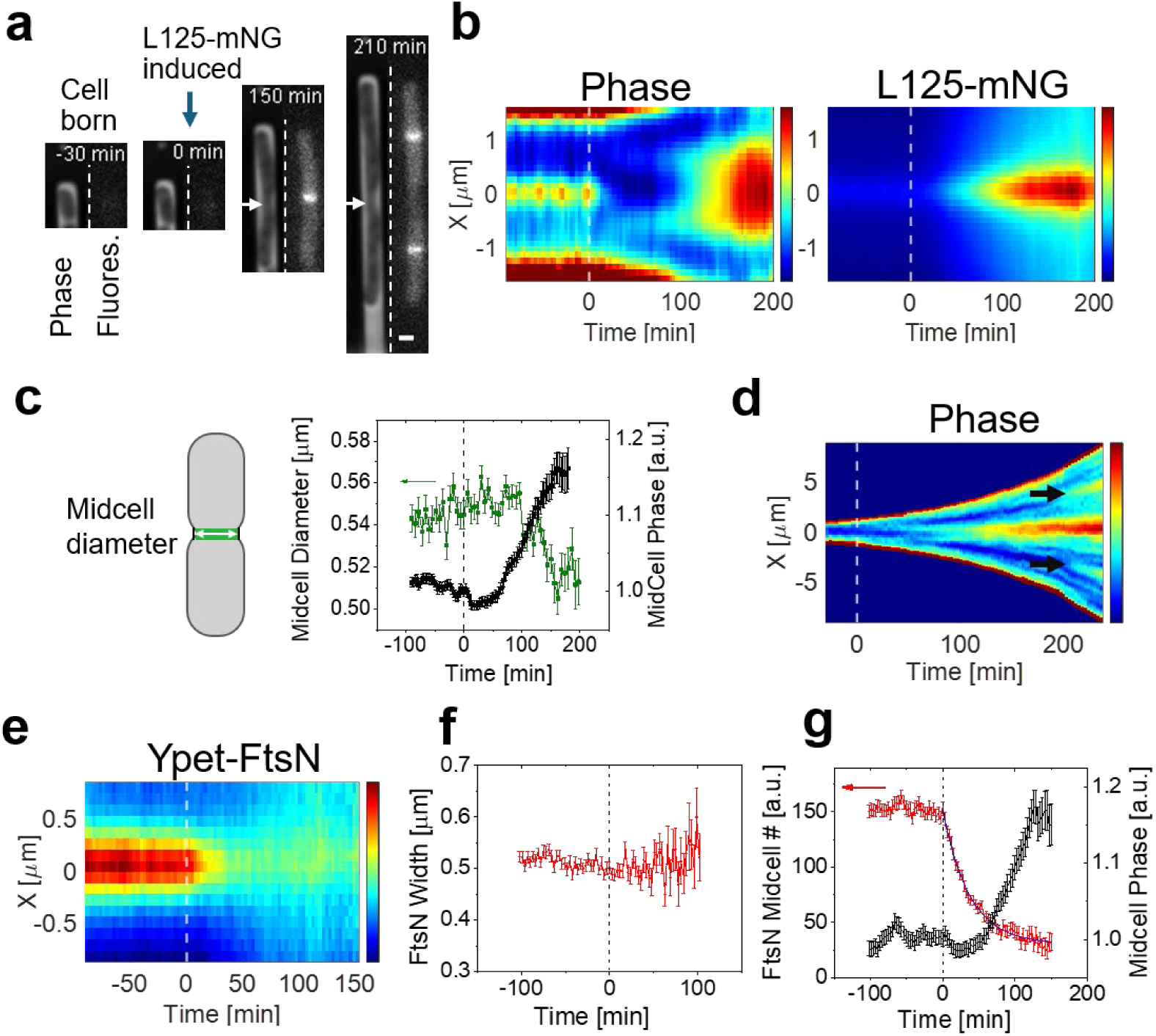
L125/L125-mNG leads to the formation of broad, shallow constrictions despite loss of the late divisome localization. **a** Representative time-lapse phase-contrast images of a cell before and after induction of L125-mNG with 30 μM IPTG. The blue arrow indicates the start of L125-mNG induction (*t* = 0 min), and white horizontal arrows indicate broad, shallow constrictions. Scale bar, 1 µm. **b** Population-averaged kymographs of phase contrast intensity (left) and L125-mNG fluorescence (right) with L125-mNG induction at *t* = 0. **c** Midcell diameter (green, left axis) and midcell phase-contrast intensity (black, right axis) as a function of time relative to L125-mNG induction. **d** Representative phase kymograph of a single cell after induction of L125-mNG. Black arrows point to constrictions forming at cell ¼ positions. **e** Kymograph of Ypet-FtsN signal before and after induction of L125 with 30 μM IPTG. Only the central region of the cells, extending ±0.8 µm from midcell, is shown. **f** Width of the midcell Ypet-FtsN accumulation as a function of time relative to L125 induction. **g** Ypet-FtsN numbers at the Z-ring as a function of time relative to L125 induction (left axis). The numbers decay exponentially after induction of the binder with a characteristic time of 37 min. Midcell phase contrast intensity as a function of time for the same cells (right axis). For panels b-d *N* = 2127 (strain JM344); for e-g *N* = 731 (strain JM333). Error bars are s.e.m.

Interestingly, the phase-contrast signal showed that the constriction zone in L125-mNG-expressing cells extended over a substantially greater distance along the cell axis than the L125-mNG accumulation zone (Fig. 4b, left versus right). In shorter cells, the phase signal along the long axis appeared as a doublet, likely reflecting phase-contrast overshoot at the edges of a single broad constriction (Supplementary Fig. 5a,b). The two peaks were separated by approximately 1.2 µm, nearly twice the width of the L125-mNG midcell accumulation (∼0.66 µm) (Supplementary Fig. 5c). In longer cells, the two peaks in phase contrast signal merged, and the resulting maximum coincided with the L125-mNG maximum, spatially linking the broad constriction to the Z-ring. The appearance of broad constrictions at the cell quarter positions following relocation of the Z-ring to these sites further supports this association.

To understand how broad constrictions could form despite the loss of late divisome localization, we considered the possibility that late divisome proteins were initially recruited to the Z-ring but subsequently dispersed over a broader region when FtsA oligomerization was perturbed. Such a scenario could be consistent with the proposal that antiparallel FtsA filaments act as a rudder for the core divisome complex ^28^. Spatially dispersed divisome complexes could produce broad, diffuse fluorescence signals that we might have missed in the static images shown in Fig. 3b. To test this possibility, we performed time-lapse imaging of cells carrying endogenous Ypet-FtsN together with inducible L125. Following L125 induction, the midcell Ypet-FtsN signal disappeared rather than broadened (Fig. 4e,f), in contrast to L125-mNG, which showed some broadening after induction (Supplementary Fig. 5c). The Ypet-FtsN midcell signal decayed exponentially with a characteristic time of 37 min (Fig. 4g). In contrast, the phase signal reporting constriction began to increase only after most FtsN had left midcell (Fig. 4g). FtsK-mNG behaved similarly: its midcell signal disappeared without discernible broadening following L125 induction (Supplementary Fig. 6). These observations argue against broad constrictions arising from spatially dispersed late divisome complexes. Instead, broad constrictions can develop after midcell localization of late divisome proteins has been lost, while remaining spatially associated with the Z-ring.

### ZipA contributes to broad constrictions independently of FtsI activity

We next asked whether the broad constrictions could instead arise from preseptal PG synthesis ^32,33^. Previous studies showed that when core divisome function is inhibited, for example, with the FtsI/PBP3 inhibitor aztreonam, excess PG synthesis can still occur near midcell through a pathway involving PBP1A/PBP1B and ZipA/FtsN, with largely redundant roles within each pair ^32,33^. However, these studies did not report visible constrictions.

To test ZipA dependence, we used a *zipA* deletion strain carrying the suppressor mutation *ftsA\** (R286W), which bypasses the essential requirement for ZipA ^30,39^. Both *ftsA\** and *ΔzipA ftsA\** cells filamented after L125 or L125-mNG expression, but we observed broad constrictions only in *ftsA\** cells (Fig. 5a,b). Time-lapse imaging confirmed the absence of detectable midcell constrictions in *ΔzipA ftsA\** cells (Fig. 5c,d), although L125-mNG still localized to midcell (Supplementary Fig. 7a). Thus, formation of the broad constrictions is dependent on ZipA.

**Fig. 5.**
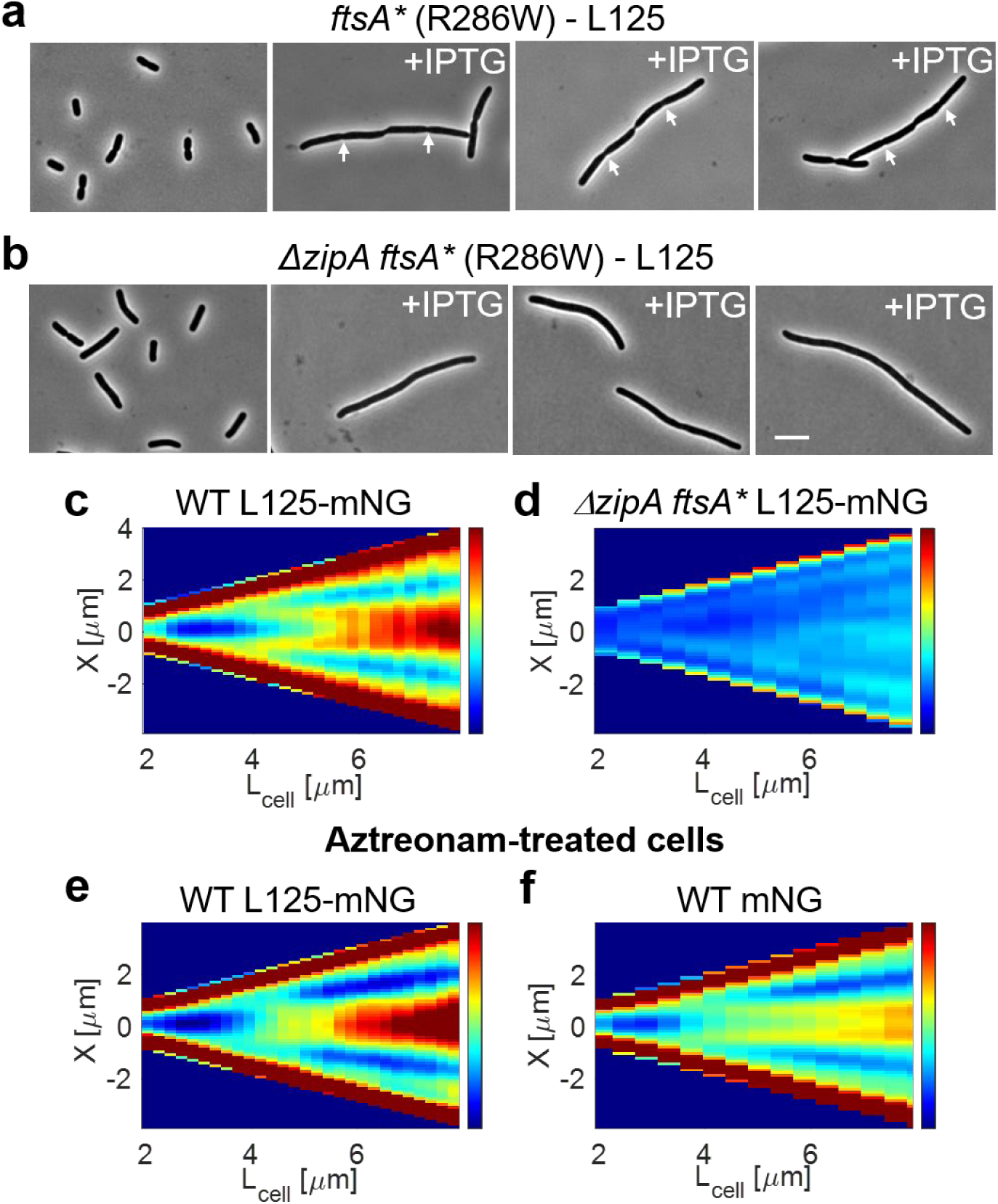
ZipA, but not FtsI activity, is required for the broad constriction phenotype. **a** Representative phase-contrast images of *E. coli ftsA\** (R286W) cells (strain JM342) before L125 induction (left) and 3 h after induction with 30 µM IPTG (right). White arrows indicate broad, shallow constrictions. **b** Representative phase-contrast images of *ΔzipA ftsA\** cells (strain JM343) under the same conditions as in panel a. Scale bar, 5 µm (shown in b; applies to a,b). **c** Cell-length kymograph of phase contrast intensity in wild-type cells expressing L125-mNG (strain JM344, *N* = 945). Cells are binned by cell length, and only cells born after the start of L125-mNG induction are included. **d** As in panel c, for Δ*zipA ftsA*\* cells expressing L125-mNG (strain JM375, *N* = 312). The weak observable patterns correspond to nucleoids. **e** As in panel c, for wild-type cells expressing L125-mNG (strain JM377, *N*=368) during simultaneous L125-mNG induction and treatment with 0.3 µg/ml aztreonam. **f** Cell-length kymograph of phase contrast intensity for control cells (strain JM228, *N*=1022) treated with 0.3 µg/ml aztreonam only. The strain does not express L125-mNG but carries a reporter plasmid expressing cytosolic mNG.

It has been reported that preseptal PG synthesis can alternatively occur through a ZipA-independent pathway involving FtsN^33^. This pathway, however, is not expected to be active in our measurements because, as explained earlier, L125-mNG prevents recruitment of FtsN to the Z-ring (Fig. 4e,f).

We next inhibited FtsI/PBP3 activity with 0.3 µg/ml aztreonam, the concentration used in earlier works ^32,33^, while simultaneously inducing L125-mNG or the mNG control. To exclude possible aztreonam degradation by plasmid-encoded β-lactamase, the ampicillin-resistance marker on the L125-mNG plasmid was replaced with a kanamycin-resistance marker. Aztreonam blocked division in both strains, but broad, shallow constriction regions still formed (Fig. 5e,f) and L125-mNG continued to localize to midcell (Supplementary Fig. 7b). In L125-mNG cells, the phase contrast pattern was similar to that observed without aztreonam (Fig. 5c,e). In the mNG control, the phase-contrast doublet was more pronounced, although the actual midcell narrowing was shallower (Fig. 5f). The corresponding constricted region in the mNG control cells was also apparent from the cytosolic mNG signal (Supplementary Fig. 7c). A possible explanation for the difference between the two aztreonam-treated strains is that L125/L125-mNG expression increases the midcell accumulation of ZipA and other early divisome components (cf. Fig. 3d), potentially promoting preseptal PG synthesis relative to aztreonam treatment alone. Regardless of this interpretation, the persistence of constrictions during FtsI inhibition, together with their loss in *ΔzipA ftsA\** cells, shows that the broad constriction phenotype requires ZipA but does not require normal FtsI/PBP3-dependent septal PG synthesis.

## Discussion

We designed a 125-residue protein binder, L125, predicted to bind the longitudinal oligomerization interface of FtsA. L125 inhibits cell division, preserves the localization of early divisome components, and blocks the recruitment of all tested essential late divisome proteins. Thus, our data support a model in which the FtsA longitudinal interface, which L125 blocks, is required for midcell recruitment of the core divisome complex.

There are at least two possibilities for how this interface can be involved in recruiting the late divisome complex. First, L125 is predicted to block the assembly of FtsA filaments and thereby also the formation of FtsA antiparallel filaments. Antiparallel filament conformation might be needed for the recruitment of late divisome proteins, which may need to contact multiple FtsA monomers in their straight conformation rather than in a curved one. Second, one could consider a scenario in which L125 sterically interferes with the binding of a late divisome protein to FtsA. FtsK is a plausible candidate because recruitment of downstream late divisome proteins depends on FtsK ^40^, and FtsK is absent from midcell after L125 induction. However, the strongly nonlinear concentration dependence of the division rate on L125-mNG, which decreases approximately exponentially above a threshold (Fig. 1f), is not readily explained by simple competition between L125-mNG and FtsK for a common binding site on FtsA. A simple one-site competitive-binding model in which division rate follows FtsK occupancy would instead be expected to yield a substantially weaker, hyperbolic, saturating dependence on the L125-mNG concentration. The observed threshold-like response is therefore more readily explained by disruption of a higher-order FtsA assembly.

We found that the cells retained slow constriction activity despite the absence of detectable late-divisome recruitment. Although originating from the Z-ring, the activity extended ∼0.6 µm to both sides of it and required ZipA. The persistence of broad constrictions during aztreonam treatment further indicates that they do not depend on normal FtsI-mediated septal PG synthesis. Together, the ZipA dependence and FtsI independence connect the phenotype to the previously described preseptal pathway ^32,33^. Broad constrictions may have escaped detection in previous studies because they progress slowly and, in static images, can be difficult to distinguish from constrictions already present when division is inhibited.

The exact function of preseptal synthesis remains to be elucidated. Its presence, however, could help explain why the Z-ring forms well before activation of the core divisome complex and why two membrane anchors, FtsA and ZipA, participate in linking FtsZ to the cell envelope. One possible function of preseptal PG synthesis is to preferentially reinforce the PG over a broad midcell region before activation of the core divisome. PG synthesis and remodeling in this region could also account for the reduction in cell diameter we observe. Such a geometry may itself confer a mechanical advantage: under turgor pressure, circumferential stress decreases as cell radius decreases, and a locally thicker PG layer would further strengthen the constriction region. Consistent with this possibility, increased PG thickness at the constriction region has been observed during early stages of constriction, when the core divisome complex was already involved ^41^. Furthermore, PBP1B, one of the enzymes implicated in preseptal synthesis, has recently been shown to fortify the division site against osmotic rupture rather than directly drive septation ^42^.

A notable feature of the broad constrictions is their approximately constant axial extent (∼1.2 μm), independent of cell length. One possible explanation is that cell wall remodeling complexes initially assemble within the narrow Z-ring and subsequently diffuse away while remaining active. If their active lifetime is limited, the distance they travel before inactivation or dissociation could define a characteristic remodeling zone whose extent is independent of cell length, provided the cell is sufficiently long. However, ZipA does not form comparably broad bands. This observation is consistent with a model in which ZipA recruits or activates the remodeling complexes at the Z-ring but does not remain associated with them during subsequent remodeling.

Developing *de novo* binders using generative machine learning holds great promise for identifying regulatory mechanisms governing cellular pathways and advancing therapeutic discovery. The usefulness of *de novo* binders in elucidating cellular pathways could exceed that of mutational screens which have been a prevailing method so far. However, the approach is not yet fully mature. A challenge we encountered in this work was developing functional binders using predefined hotspots. Although three hotspot-targeted BindCraft runs produced candidates with favorable computational metrics, none of the three candidates selected for experimental testing showed activity in cells, suggesting that such metrics alone may be insufficient to identify functional binders. In contrast, removing predefined hotspots yielded multiple binders clustered near the longitudinal interface, including L125. One possible limitation is that AlphaFold2, used during BindCraft sequence optimization, was trained before the structure of antiparallel FtsA filaments became available. Future structure-prediction and design models may make such interfaces more tractable.

FtsA is broadly conserved but has not yet been exploited as an antibiotic target ^1,43^. By contrast, small-molecule and peptide binders that alter FtsZ oligomerization already exist ^44,45^. The inhibition of cell division by L125 suggests that the FtsA oligomerization interface could likewise be targeted by small molecules or cyclic peptides.

## Methods

### Design of protein binders targeting FtsA–FtsA contacts at lateral and longitudinal interfaces

*De novo* protein binders aimed at disrupting FtsA–FtsA contacts were designed using BindCraft version 1.5 ^31^ with a 2.8 Å resolution X-ray crystal structure of *E. coli* FtsA (PDB 7Q6D ^28^) as the target. An *E. coli* FtsA double-filament model, based on the antiparallel double-filament structure of *Vibrio maritimus* FtsA reported by Nierhaus et al. ^28^, was used to define FtsA–FtsA interface residues. Two design approaches were used: hotspot-targeted and hotspot-free design.

For the hotspot-targeted approach, three BindCraft runs were performed. The first used the exposed hydrophobic residues I94, M96, V97, V119, V121, I143, V147, L149, and V152 as hotspot residues. Binders were generated with lengths between 70 and 150 residues. The second used I94, M96, V97, Y139, and V152 as hotspots. To reduce computational cost, a trimmed version of FtsA was used that included only residues 87 to 331. Binders ranged from 50 to 80 residues in length. The third used I137, I143, and L149 as hotspots, and the complete FtsA structure as the target. Each run produced designs that were filtered based on default BindCraft criteria (Supplementary Data 1, Hotspot-targeted). Although no designs passed all the default filtering criteria, three candidates targeting the lateral interface were selected for experimental testing based on high interface predicted TM scores (ipTM>0.85) and compactness (Supplementary Table 1). The corresponding coding sequences were cloned into inducible expression constructs as described below.

In a second design strategy, the hotspot constraint was removed, allowing BindCraft to generate FtsA binders ranging from 60 to 180 amino acids in length without any predefined interface requirements. The untrimmed FtsA X-ray structure was used as the target. This campaign yielded 18 candidates that passed the default BindCraft design filters; these proteins are listed in Supplementary Data 1, Hotspot-free designs. Four longitudinal interface binders were selected for experimental testing (Supplementary Table 1).

### Structural analysis of predicted FtsA–binder complexes

Predicted FtsA–binder complexes were aligned to an FtsA monomer within the antiparallel FtsA double-filament structure (PDB 7Q6F) using PyMOL version 3.1.0 (Schrödinger, LLC) to visualize their positions relative to the FtsA–FtsA interfaces. Intermolecular contacts between FtsA and L125 in the predicted complex were identified using the Protein–Ligand Interaction Profiler (PLIP) ^46^ with default interaction-detection parameters. Hydrogen bonds, salt bridges and hydrophobic contacts identified by PLIP were used for the structural interaction analysis shown in Supplementary Fig. 1b,c.

### Strain and plasmid construction

Strains, plasmids, oligonucleotides, and binder-coding sequences used in this study are listed in Supplementary Tables 2, 3, 4, and 5, respectively.

### Construction of inducible binder expression plasmids

Inducible expression constructs for the FtsA lateral-interface binders Lat62, Lat66 and Lat123 (Supplementary Table 5) were generated from pSEB306+ ^47^, a pDSW210-derived plasmid carrying full-length *ftsA* under the control of a weakened Ptrc promoter. Binder-coding sequences were codon-optimized for expression in *E. coli* using the IDT Codon Optimization Tool. Codon-optimized inserts containing the native *ftsA* 5ʹ untranslated region and terminal sequences homologous to the pDSW210-derived backbone were synthesized by Integrated DNA Technologies Inc. pSEB306+ was digested with NcoI and HindIII, the *ftsA* coding sequence was replaced with each binder-coding sequence using the NEBuilder® HiFi DNA Assembly Kit (NEB). Plasmids were propagated in DH5α with ampicillin selection at 100 μg/ml. The same cloning strategy was used to generate inducible constructs expressing the longitudinal-interface binders L75, L81, L102 and L125.

C-terminal mNG fusions to the FtsA protein binder candidates were generated using NEBuilder® HiFi DNA Assembly. Vector backbones were amplified from the corresponding binder-expression plasmids, and the mNG coding sequence was amplified from pJM21 using Q5® Hot Start High-Fidelity DNA Polymerase (NEB). The binder stop codon and mNG start codon were omitted to create in-frame binder–mNG fusions, and an EcoRI site was introduced at the binder–mNG junction. Assembly primers were designed using the NEBuilder® Assembly Tool and are listed in Supplementary Table 4.

To eliminate plasmid-encoded β-lactamase activity as a potential confounding factor in the aztreonam treatment experiments, the bla/Amp^R^ marker of pJM-L125-mNG was replaced with an aph/Kan^R^ cassette amplified from pJM21. Plasmids used and constructed in this study are listed in Supplementary Table 3. Constructs were verified by restriction analysis and either whole-plasmid sequencing (Plasmidsaurus) or Sanger sequencing of the insert and assembly junctions (Eurofins Genomics).

### Construction of chromosomal fluorescent fusions

Chromosomal fluorescent fusions to the divisome proteins FtsK, FtsX, and FtsI were generated at the native chromosomal loci by λ Red recombineering using established approaches ^48, 49^. FtsK and FtsX were tagged at their C-termini to generate FtsK–mNG and FtsX–mNG, whereas FtsI was tagged at its N-terminus to generate mNG–FtsI. Recombineering fragments were amplified using Q5® Hot Start High-Fidelity DNA Polymerase and primers containing homology to the corresponding chromosomal target regions.

For the N-terminal mNG–FtsI fusion, BW25113 cells carrying pKD46 were electroporated with a PCR product amplified from pJM139, which contains an FRT-*aph*-FRT-mNG-linker cassette encoding the SAGSAAGSGSA linker. The recombineering fragment was amplified using primers PR229 and PR230. For the C-terminal FtsK–mNG and FtsX–mNG fusions, recombineering fragments were amplified from pJM21, which contains an SSAGSAAGSGEF linker–mNG–FRT-*aph*-FRT cassette, using primers PR233/PR234 and PR231/PR232, respectively. These fragments were electroporated into JM217 cells carrying pSIM5, which encodes the λ Red recombination proteins.

PCR products were treated with DpnI, purified using the Monarch® DNA Gel Extraction Kit or Monarch® PCR & DNA Cleanup Kit, and electroporated into the appropriate recombineering host strain. Recombinant colonies were selected on LB agar containing kanamycin at 25 μg/ml. Correct chromosomal insertion was confirmed by colony PCR, and fluorescent signal and divisome localization were verified by fluorescence microscopy. Oligonucleotides used for recombineering are listed in Supplementary Table 4, and strains used and constructed in this study are listed in Supplementary Table 2.

### Growth media and growth conditions

For *E. coli* strain construction, cells were grown in Luria-Bertani (LB) broth supplemented with appropriate selective antibiotics. For time-lapse imaging in microfluidic devices, cells were cultured at 28°C in M9 minimal medium supplemented with 2 mM magnesium sulfate, 0.5% glucose, 0.5% casamino acids and 1× trace metals mixture. When required, medium was supplemented with ampicillin at 100 μg/ml or kanamycin at 40 μg/ml. For induction, IPTG was added to the medium at 30 μM or 100 μM.

### Cell preparation, binder induction and aztreonam treatment in microfluidic devices

Strains were streaked on M9 minimal medium agar plates supplemented with 2 mM MgSO₄, 0.5% glucose, 0.5% casamino acids, 1× trace-metal solution and the appropriate selective antibiotics. For microscopy experiments, single colonies from freshly streaked plates were inoculated into 3 ml of liquid M9 medium supplemented with the same components and grown to early exponential phase (OD₆₀₀ ≈ 0.1). Strains carrying protein-expression plasmids were grown with ampicillin at 100 µg/ml, except for constructs in which the ampicillin-resistance marker had been replaced with kanamycin resistance; these strains were grown with kanamycin at 40 µg/ml.

Bovine serum albumin (BSA; MilliporeSigma, MO, USA) was added to the cultures to a final concentration of 0.075 mg/ml to reduce cell clumping. Cells were then concentrated approximately 100-fold by centrifugation and loaded into PDMS-based mother-machine microfluidic devices fabricated as described previously ^50^. When indicated, two or three strains were mixed before loading so that they could be cultured and imaged on the same microfluidic device, with one of the strains serving as a wild-type reference and the other carrying a fluorescent reporter for comparison. These strains were later distinguished by their fluorescent signatures/

The concentrated cell suspension was pipetted into the main flow channel of the device and allowed to populate the dead-end growth channels for 1 h. Fresh M9 growth medium containing the appropriate carbon source, additives, BSA and selective antibiotics, when required, was supplied from a 10-ml syringe mounted on an NE-1000 syringe pump (New Era Pump Systems, NY, USA). Medium flow was maintained at 6 µl min⁻¹ throughout the experiment. Cells were grown in the microfluidic channels for at least 14 h before imaging to establish steady-state growth and to ensure that by the start of imaging, the populated channels contained descendants of a single initially trapped cell.

For binder induction and binder-induction/aztreonam experiments, two 10-ml syringes were prepared the following morning: one containing fresh M9 growth medium without inducer and one containing otherwise identical medium supplemented with 30 µM IPTG. For aztreonam experiments, aztreonam was included in the IPTG-containing medium at 0.3 µg/ml. The syringes were mounted on two separate NE-1000 syringe pumps and connected to the microfluidic device through a T-junction connector. After 10 h of imaging in non-inducing medium, expression of the FtsA protein binder and the mNG reporter was induced by switching the flow to IPTG-containing medium. Medium exchange was performed without interrupting image acquisition by turning off the pump containing non-inducing medium and turning on the pump containing IPTG-supplemented medium. In aztreonam experiments, IPTG-supplemented medium also contained aztreonam. Syringe pumps were controlled using a custom LabVIEW program. All microfluidic imaging experiments were performed at 28 °C and lasted typically 22–26 h.

### Cell fixation for static images

Cells were grown overnight at 28 °C in M9 medium supplemented with 0.5% glucose and 0.5% casamino acids (M9 GluCas). The following morning, cultures were diluted into 3 ml fresh M9 GluCas medium and subsequently diluted again into 3 ml fresh medium in the afternoon to maintain exponential growth. When cultures reached an OD₆₀₀ of approximately 0.05–0.06, 1 ml aliquots were either left uninduced or induced with 30 µM IPTG for 180 min. Uninduced and induced samples, with or without pJM-L125, were fixed in parallel at the same time. Cells were fixed by adding paraformaldehyde and glutaraldehyde directly to the culture to final concentrations of 2.8% and 0.04%, respectively, and then incubated for 10 min at room temperature. Fixed cells were washed with 10% glycerol and stored at 4 °C until imaging. For static imaging, fixed cells were immobilized on 2% agarose pads and covered with a coverslip for microscopy.

### Fluorescence microscopy

Bacteria were imaged using a Nikon Ti-E inverted fluorescence microscope (Nikon Instruments, Japan) equipped with a 100× NA 1.40 oil-immersion phase-contrast objective, an iXon DU897 EMCCD camera (Andor Technology, Ireland), a motorized stage (Prior Scientific, MA, USA), and the Nikon Perfect Focus® system for time-lapse imaging. Image acquisition was controlled using NIS-Elements software (Nikon Instruments). Fluorophores were excited with a 200 W Hg lamp. For time-lapse imaging, excitation light was attenuated using ND4 and ND8 neutral-density filters. Chroma 41001 and 41004 filter sets (Chroma Technology Corp., VT, USA) were used to image mNG/Ypet/sfGFP and mCherry, respectively.

Time-lapse imaging in M9 glucose-casamino acid medium was performed at 3-min intervals. Typical exposure conditions were 400 ms with an EM gain of 200 for mNG/Ypet/sfGFP fluorescence, 200 ms with an EM gain of 200 for mCherry, and 400 ms without EM gain for phase-contrast images.

For static fluorescence imaging of fixed cells, mNG, Ypet and mCherry images were acquired with a 1 s exposure and an EM gain of 200, and phase-contrast images were acquired with a 1 s exposure without EM gain. Neutral-density filters were removed for static imaging.

### Image analysis

MATLAB, along with the Image Analysis Toolbox and DipImage Toolbox (http://www.diplib.org/), was used for image analysis. In all analyses of time-lapse recordings, corrections to subpixel shifts between different frames were applied first. These shifts were determined by correlating phase-contrast images in adjacent frames. The cells were then simultaneously segmented and sorted into lineages based on phase-contrast images using a custom MATLAB script. The script tracked the peaks in phase contrast intensity along the length of the channel. The peaks in the phase contrast intensity profile correspond to cell poles.

The cell coordinates from the lineages were used to extract intensity line profiles of fluorescent and phase images along the long axis of the cells. For the intensity line profiles, 11 individual profiles covering a band about 1.2 µm wide centered on the channel’s symmetry axis were averaged. For a coarse background correction for each cell, the average intensity from an equally wide band, but shifted 1.8 µm from the channel, was subtracted. The procedure yielded the intensity line profiles *I*_0_(X, *t*) for each cell along its long-axis coordinate X at time *t*. *t* = 0 in all measurements corresponds to the treatment start time. To correct for cell autofluorescence, we also imaged an unlabeled reference strain (BW27783 or JM320) on the same microfluidic chip and determined *I*_0,W*T*_(X, *t*) for these cells too. We then calculated the average per pixel intensity, *I_bg_* = ∑*_X_ I*_0,W*T*_(X, *t*) /*l_cell_* from these cells, averaging over all measurements of all cells. Here *l_cell_* is cell length in pixels. The corrected fluorescence intensity used in measurements is then *I*(X, *t*) = *I*_0_(X, *t*) – *I_bg_*. Note that no background was subtracted from the average line profiles for the phase images.

### Determination of L125 concentration

L125 concentrations were determined based on fluorescence measurements of its fusion to mNG (L125-mNG; strain JM344) or from fluorescence measurements of mNG (strain JM298). In both measurements, L125-mNG or mNG was expressed from the same plasmid with the same promoter and ribosome binding site as L125.

We first calculated the total fluorescence intensity from the cell at each recorded time *I_tot_*(*t*) = ∑*_X_ I*(X, *t*). To obtain uncalibrated concentration in steady state conditions, the background-subtracted total intensity was divided by the cell length, *l_cell_* that is *i*(*t*) = *I_tot_*(*t*)/*l_cell_*(*t*). This quantity is plotted in Fig. 1c.

At early times after induction of L125-mNG, correction for fluorophore maturation is needed to accurately capture its total concentration, which includes L125 molecules with both mature and immature (not yet fluorescent) fluorophores. Following the approach outlined in the earlier study ^19^, the correction factor is calculated based on the expected concentrations of mature (*i_m_*) and the total protein (*i_tot_*) expressed via fluorescent intensities per unit length of the cell for *t* > 0 as:

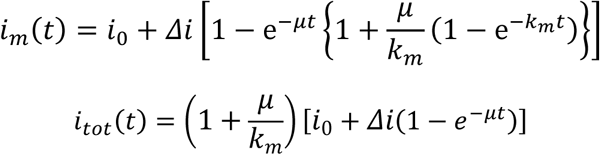

Here *μ* is the growth rate determined for each cell separately based on its elongation rate and *k_m_* is the maturation rate for mNG. In our growth conditions *k_m_* = 0.029 *min*^−1^ ^19^ taken the same for all cells. *i*_0_ is the average uncalibrated concentration of cells in steady-state conditions before induction and *Δi* the total increase of the intensity as the cells reach a new steady-state after induction. These values of *i*_0_ and *Δi* are found from the fit of *i_m_*(*t*) to data as shown in Fig. 1c. The correction fraction, *F*(*t*), is then expressed as:

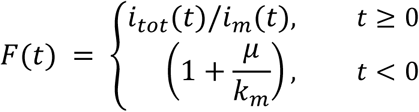

This correction factor is applied on a single-cell basis, yielding fluorescence maturation corrected concentration, which, however, is not yet calibrated:

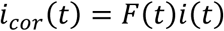

We expressed concentrations of L125-mNG, [*L*125*mNG*], in terms of estimated FtsA concentration in the cell [*FtsA*] that is [*L*125*mNG*]/[*FtsA*]. To find the conversion from uncalibrated concentration, *i_cor_*, to [*L*125*mNG*]/[*FtsA*] we determined [*L*125*mNG*] relative to FtsZ concentration in the cell, [*FtsZ*]. We then used published mass spectrometry data for ([*FtsZ*]/[*FtsA*])*_m_*_.*s*._to find the conversion coefficient between *i_cor_* and [*L*125*mNG*]/[*FtsA*]. Note that we used fluorescent fusion to FtsZ, even though we were interested in the L125-mNG concentration relative to that of FtsA, because no equivalent strain with FtsA genomic fusion is available. The time-dependent concentration is then expressed as:

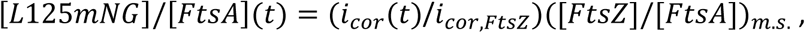

where *i_cor_*_,*FtsZ*_ is an uncalibrated (constant) concentration for fluorescent fusion to FtsZ. To find *i_cor_*_,*FtsZ*_ we used strain JM147 in which the native FtsZ is replaced with an FtsZ sandwich fusion to mNG ^51^. Previous Western blot analysis shows that JM147 expresses FtsZ-mNG at the level of WT FtsZ (strain BW27783) ^19^. To find the corresponding *i_cor_* we used mNG reporter strain JM298. We chose to use mNG reporter instead of L125-mNG because only a few L125-mNG cells could be grown to steady expression levels in microfluidic channels, and these cells expressed less than an average amount of L125-mNG. We mixed JM147 along with JM298 expressing cytosolic mNG and the WT strain BW27783, and carried out a time-lapse measurement of this strain mixture on a mother machine device. We determined the uncalibrated concentrations of FtsZ-mNG, *i_cor_*_,*FtsZ*_, and mNG, *i_cor_*_,*mNG*_(*t*), from this measurement as explained above and calculated their ratio (Supplementary Fig. 2a). Fitting this ratio to the equation for *i_m_*(*t*) yielded a steady state value *i_cor_*_,*steady*_/*i_cor_*_,*FtsZ*,*steady*_ ≈ 0.25 at 30 µM IPTG induction. In terms of these steady state concentrations [*L*125*mNG*]/[*FtsA*](*t*) can be expressed as:

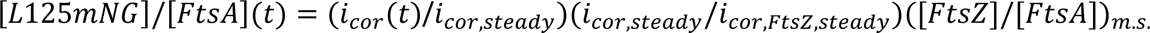

For ([*FtsZ*]/[*FtsA*])*_m_*_.*s*._we used a comprehensive compilation of mass spectrometry data for *E. coli* grown in a glucose-casamino acid medium from different groups ^52^. Across the compiled datasets, most measurements yielded ([*FtsZ*]/[*FtsA*])*_m_*_.*s*._ ≈ 5…6. Based on these data we chose 5.5 as the representative value. This choice yielded ([*L*125*mNG*]/[*FtsA*])*_steady_* = 5.5 · 0.25 = 1.4 at steady state when induced with 30 µM IPTG. For L125-mNG for 30 µM IPTG induction *i_cor_*_,*steady*_ = 346. This yielded the final calibration factor (1/*i_cor_*_,*steady*_)(*i_cor_*_,*steady*_/*i_cor_*_,*FtsZ*,*steady*_)([*FtsZ*]/[*FtsA*])*_m_*_.*s*._= 0.004. Thus, ([*L*125*mNG*]/[*FtsA*])(*t*) = 0.0040 *i_cor_*(*t*).

### Determining the average cell population division rate

The population-average division rate was determined as a function of [L125-mNG]/[FtsA] (Fig. 1f). [L125-mNG]/[FtsA] was determined at each cell division, and the resulting events were binned by this concentration. For each concentration bin, we counted the number of division events, *N_div_*.

To determine the division rate, we also calculated the total number of eligible cell-frames, *N_frame_,* contributing to each concentration bin. For this calculation, [L125-mNG]/[FtsA] was determined for every recorded cell-frame and binned using the same concentration-bin edges as for the division events. The population-average division rate in each bin was then calculated as

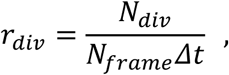

where Δ*t* is the time between consecutive frames.

Error bars for the division rates were calculated, assuming Poisson statistics for the number of observed events in each concentration bin. Exact 68% Garwood Poisson confidence intervals for the event counts were calculated and converted to confidence intervals for the corresponding rates by dividing the lower and upper count limits by the total cell-time exposure, *N*_frame_Δ*t*.

### Kymographs based on cell length, cell age and aligned relative to treatment start time

**Cell-length kymographs** were generated by binning individual-cell intensity profiles by cell length. Cell lengths were grouped into bins of an integer number of pixels. For each cell-length bin, intensity profiles from individual cells were aligned at midcell and placed in a common spatial coordinate system by symmetrically zero-padding the profiles to the maximum cell length included in the analysis. Profiles were oriented such that the old pole was on the negative side of the spatial coordinate X. To prevent slowly elongating cells from being overrepresented, each individual cell was allowed to contribute at most one intensity profile to a given cell-length bin. Profiles within each bin were then summed and divided by the number of contributing cells to obtain the population-averaged intensity profile.

**Cell age kymographs** were created by averaging *I*(X, *t*) from individual cells at fixed cell cycle ages over the cell population. The cell age, *a*, is calculated as *a* = *t*/*T_d_* where *T_d_* is the duration of the cell cycle and *t* is the time from the cell’s birth. Since cells at a given cell age are of different lengths, the cell coordinates, X, are scaled by the average length of an elongation curve. For the division cycle kymographs, the average length of the elongation curve is *L_cell_*_,*av*_(*a*) =< *L_cell_*_,*b*_ > 2*^a^* where < *L_cell_*_,*b*_ > is the population average cell length at birth as determined from the same measurement. The new coordinate values for each cell, *i*, at the age *a* are X^′^ = X *L_cell_*_,*av*_(*a*)/*L_cell_*_,*i*_(*a*) where *L_cell_*_,*i*_(*a*) is the length of this cell at age *a*. The *I*(X^′^, *a*) values are then resampled at a fixed spatial grid and the resulting curves are averaged over different cells. The origin of the coordinate axis is chosen at the center of the cell. The most negative coordinate value corresponds to the old pole of the cell.

**For treatment-aligned kymographs,** fluorescence intensity profiles from different cells were averaged in each measurement frame around the L125 induction time (*t* = 0 in all graphs). In these kymographs, cell midpoints were aligned, and the intensity profiles of cells of different lengths were averaged at a given time point. Kymograph presents only a limited region (X = ±0.8 μm for Fig. 4e) from the cell middle.

### Determining midcell protein excess amounts and excess fractions

To determine midcell protein excess amounts and excess fractions, the integrated signal intensity of corresponding fluorescence labels in a 0.75 μm wide band centered at the midcell was calculated using:

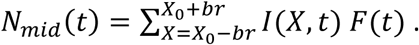

Here X_0_ and *br* denote the cell center coordinate and half-width of the band, respectively. *br* was taken 3 pixels so that the total width of the band was 7 pixels (0.75 µm). Similarly, the cell background value was calculated at the cell ¼ coordinate X_0.25_ as

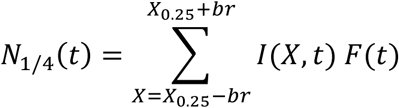

The excess midcell amount is calculated as:

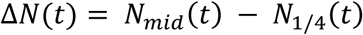

The excess amount for a given protein is proportional to the number of these proteins in the Z-ring. However, this number has not been calibrated and is given in arbitrary units. The midcell amount curves shown in the same plot have the same arbitrary-unit scales, but the arbitrary-unit scales for different labeled proteins are not comparable.

The relative excess midcell fraction is calculated as:

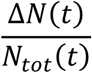

*N_tot_*(*t*) = ∑ *I*(X, *t*)*F*(*t*) is the estimated total number of proteins in the cell. The excess midcell amount and the excess midcell fraction are referred to as the midcell amount and the midcell fraction, for brevity.

### Determining the width of midcell accumulations

To determine the width of midcell accumulations of fluorescently labeled proteins, fluorescence intensity line profiles *I*(X, *t*) were fitted to a Gaussian function plus a constant background term. Four pixels adjacent to each cell pole were excluded from the fit. The fit parameters included the position and amplitude of the fluorescence maximum, the full width at half maximum (FWHM) of the Gaussian, and the constant background intensity. Fits were excluded if any of the following criteria were met: 1) the fitted position of the fluorescence maximum was more than 8% of the cell length from the cell center, 2) FWHM < 0.2 µm, or 3) FWHM > 1.3 µm. 0.2 µm represents an unrealistically narrow fit relative to optical resolution, and 1.3 µm excludes too-wide ones that cover more than 2/3 of the newborn cell length. 8% offset is also unrealistic, as 68% maxima are within 2.5% of the cell center. The widths plotted in the figures correspond to the fitted FWHM.

### Determining the midcell diameter from fluorescent images

To determine the midcell diameter, fluorescence intensity profiles along the short axes of the cell were collected from a 0.75-µm-wide band and background correction was applied. The profiles were fitted to two Gaussians plus a background term. The distance between the centers of the Gaussians was used as a proxy for cell diameter. The method would not provide an adequate proxy when the cells are completing the constrictions, but L125-treated cells never reached this stage.

## Data availability

The source data generated during and/or analyzed during the current study are available from the corresponding author.

## Acknowledgments

The authors thank Harold Erickson and Sven van Teeffelen for their valuable comments, Chathuddasie Amarasinghe for help with microfluidic chip fabrication, David Weiss, Piet de Boer, Joe Lutkenhaus, and Sebastien Pichoff for bacterial strains. A part of this research was conducted at the Center for Nanophase Materials Sciences, which is a US Department of Energy Office of Science User Facility at Oak Ridge National Laboratory. This work was supported by the National Institutes of Health award GM127413 (JM) and NSF MCB2313719 (JM). JMP was supported by the Laboratory Directed Research and Development program at Oak Ridge National Laboratory, which is managed by UT-Battelle under contract DE-AC05-00OR22725 for the US Department of Energy.

## Author contributions

JaanM and JaanaM designed the experiments, JaanaM contributed all the bacterial strains and plasmids and carried out the experiments, JaanM and JaanaM analyzed the data, JMM and JMP generated the protein designs, and JaanM wrote the manuscript with input from all the authors.

## Notes

### Competing Interest Statement

The authors have declared no competing interest.

