## Supplementary Information for "A *de novo* FtsA binder blocks late divisome recruitment, unmasking a hidden constriction mechanism"

Supplementary Figures

**Supplementary Fig. 1.** Predicted binding of L125 and other *de novo* binders to the FtsA longitudinal interface.

**Supplementary Fig. 2.** Quantification of L125-mNG expression and division phenotypes after L125 induction.

**Supplementary Fig. 3.** L125-mNG localization relative to divisome proteins and during cell filamentation.

**Supplementary Fig. 4.** Broad, shallow constrictions form at midcell and cell quarter positions in L125-mNG expressing cells (JM344).

**Supplementary Fig. 5.** The broad, shallow constrictions are much wider than the midcell accumulations of L125-mNG and the early Z-ring proteins.

**Supplementary Fig. 6.** FtsK localization after L125 induction.

**Supplementary Fig. 7.** ZipA is required for the formation of L125-induced constrictions, which persist during aztreonam treatment.

Supplementary Tables

**Supplementary Table 1**. AlphaFold2 confidence metrics for de novo FtsA interface binder candidates.

**Supplementary Table 2.** Strains used in this study.

**Supplementary Table 3.** Plasmids used in the study.

**Supplementary Table 4.** Oligonucleotides used in this study.

**Supplementary Table 5.** Coding sequences of experimentally tested FtsA interface binders.

Supplementary Data 1

***De novo* protein sequences and scores as output by BindCraft from hotspot-targeted (the 1st sheet) and hotspot-free (the 2$^{nd}$ sheet).** Pink shading indicates experimentally tested designs that did not affect cell division, and green shading indicates the design that did show the effect.

Movie M1

**Growth of *E. coli* JM344 cells in the microfluidic mother machine channel before and after induction of L125-mNG expression.** The expression is induced at time 0 min. The left panels show the phase, and the right panel shows L125-mNG fluorescence images. Scale bar corresponds to 1 μm.

a

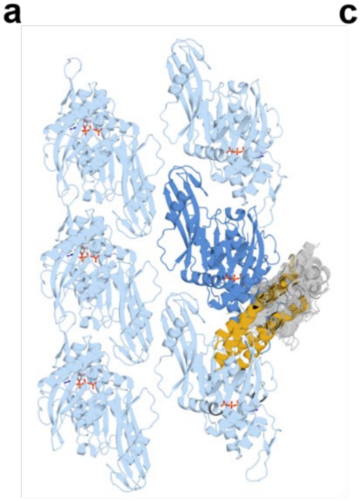

c

| FtsA | L125 contacts |
|---|---|
| Arg44 | Glu112 (HB); Glu115 (SB) |
| Lys59 | Glu115 (SB) |
| Arg63 | Ser75 (2 HB); Asn105 (HB); Tyr108 (Hyd); Glu112 (SB) |
| Asp66 | Tyr108 (Hyd, HB) |
| Gln67 | His78 (HB); Trp101 (Hyd, HB) |
| Leu70 | Phe97, Lys100, Leu104 (all Hyd) |
| Met71 | Phe97 (Hyd) |
| Pro250 | Trp55 (Hyd) |
| Ser252 | Asp51 (HB); Tyr66 (2 HB) |
| Asp253 | His52 (SB) |
| Glu255 | Trp74 (Hyd) |
| Asp369 | Lys89 (Hyd); Pro90 (HB) |
| Tyr370 | Thr85 (Hyd), Lys89 (Hyd); Phe86 (HB) |

Hyd, hydrophobic interaction; HB, hydrogen bond; SB, salt bridge

b

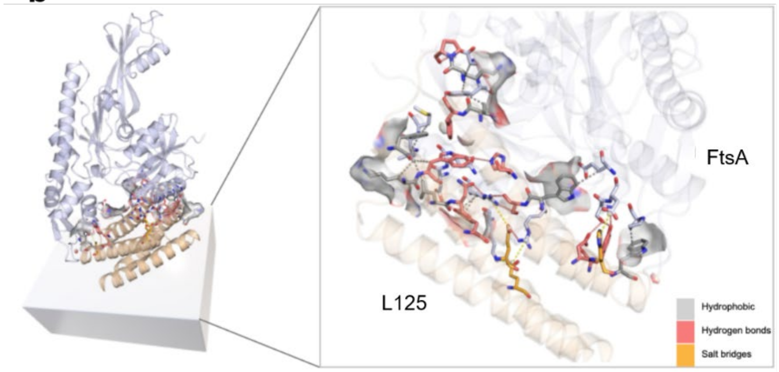


**Supplementary Fig. 1. Predicted binding of L125 and other *de novo* binders to the FtsA longitudinal interface. a** Overlay of 18 *de novo* binders designed to target the longitudinal FtsA-FtsA interface. Predicted FtsA-binder complexes were aligned to an FtsA monomer (dark blue) within the antiparallel FtsA double filament structure (light blue, PDB 7Q6F). L125 is shown in gold, and the other 17 binders are shown in light grey; three of these were also experimentally tested. **b** Predicted L125-FtsA interface. Noncovalent interactions identified by Protein-Ligand Interaction Profiler (PLIP) are shown as sticks and surfaces. FtsA is light blue, and L125 is gold. L125 residues are colored by dominant interaction type: hydrophobic interactions (grey), hydrogen bonds (red), or salt bridges (orange), with salt bridges prioritized over hydrogen bonds and hydrophobic interactions. **c** PLIP-predicted noncovalent interactions between FtsA and L125. FtsA residues shaded light green belong to domain IIB; unshaded residues belong to domain IA. HB, hydrogen bond; Hyd, hydrophobic interaction; SB, salt bridge.

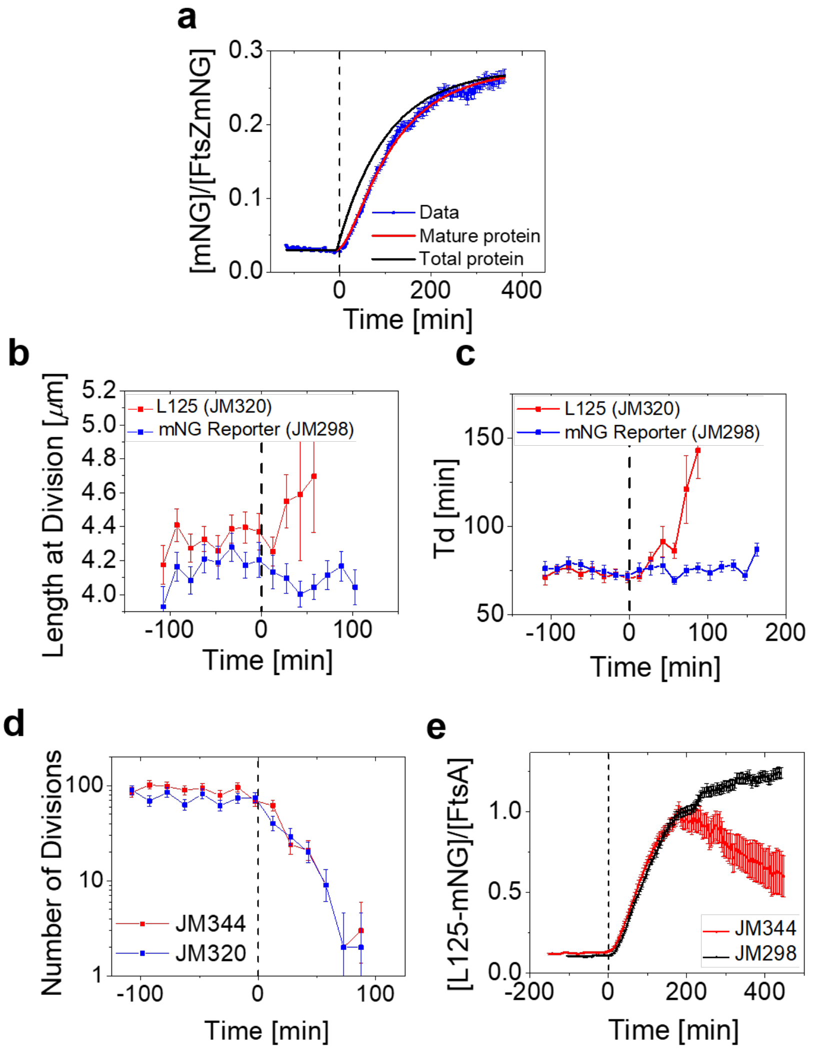


**Supplementary Fig. 2. Quantification of L125-mNG expression and division phenotypes after L125 induction. a** Ratio of mNG reporter (JM298, *N*=4343) concentration to FtsZ-mNG (JM147, *N*=1660) concentration as a function of time. FtsZ-mNG is expressed from the native *ftsZ* locus. The dashed vertical line indicates the start of L125-mNG induction (t = 0 min). The red line shows a fit to the mature [mNG] to [FtsZ-mNG] ratio, whereas the black line shows the predicted total [mNG] to [FtsZ-mNG] ratio. The baseline value from the fit is 0.030, and the total increase above baseline is 0.24. **b** Cell length at division as a function of time for strains carrying the inducible L125 plasmid (JM320, *N*=676) or the mNG reporter plasmid (JM298, *N*=815 cells). **c** Doubling time as a function of time for the same strains. **d** The number of cell divisions per 15-min interval as a function of time for strains expressing L125 (strain JM320, *N*=702) or L125-mNG (JM344, *N*=834). Both strains are from the same experiment. **e** Comparison of L125-mNG concentration inferred for JM344 with that inferred from the mNG reporter strain JM298. The data was obtained in two separate measurements, both induced with 30 µM IPTG. Error bars are s.e.m. except in panel d, where 68% Poisson confidence intervals are shown.

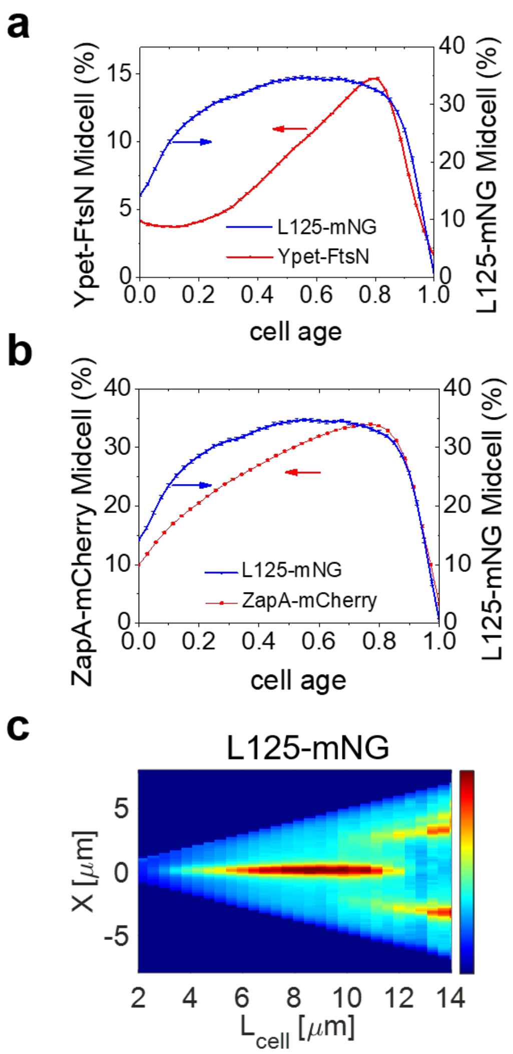


**Supplementary Fig. 3. L125-mNG localization relative to divisome proteins and during cell filamentation. a** Comparison of the midcell fractions of L125-mNG (strain JM344; *N*=3580; right axis) and Ypet-FtsN (strain JM333, *N*=5700; left axis) before L125-mNG induction, when L125-mNG was expressed at its basal level. **b** The same comparison between L125-mNG and ZapA-mCherry (strain JM333, *N*=5700). Error bars in panels a and b are s.e.m. **c** Cell length kymograph of L125-mNG fluorescence in JM344 cells after induction with 30 µM IPTG. Cells were binned by length, and the L125-mNG intensity profiles were averaged within each bin. As cells elongate to approximately 10 µm, the midcell L125-mNG band begins to split into two bands near the ¼ positions, consistent with the formation of new Z-rings at these positions.

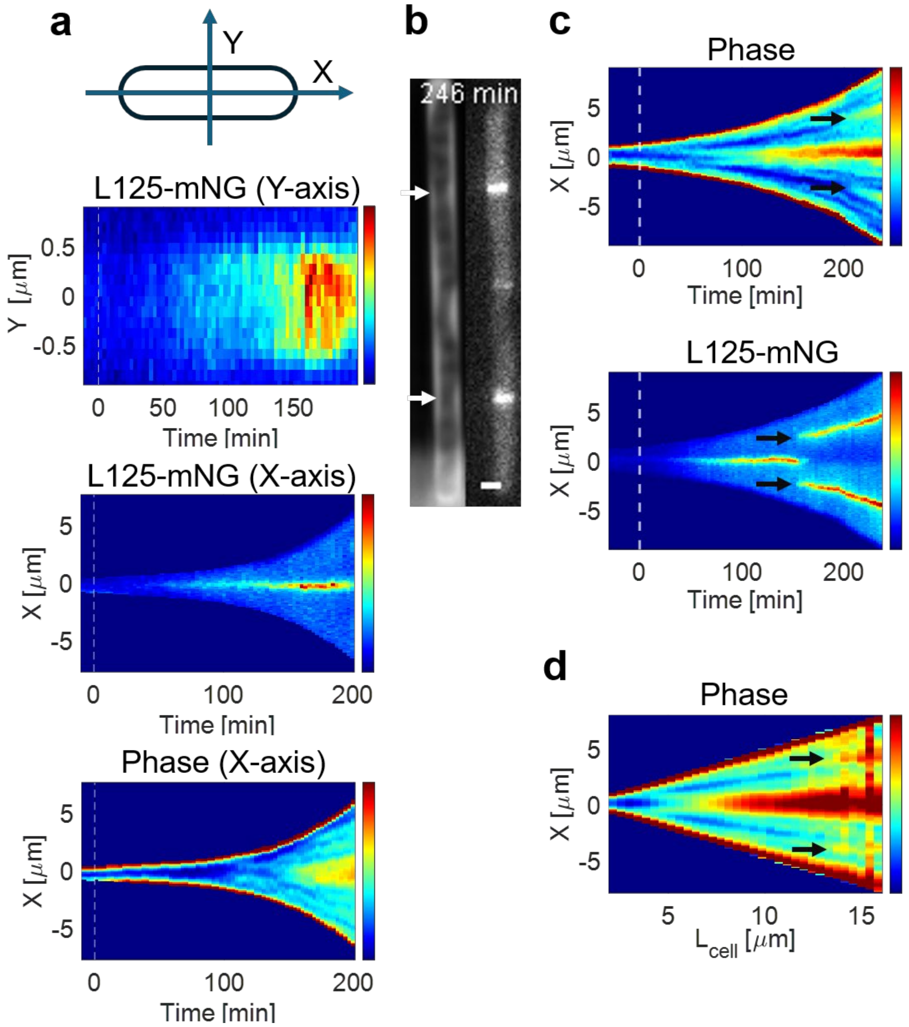


**Supplementary Fig. 4. Broad, shallow constrictions form at midcell and cell quarter positions in L125-mNG expressing cells (JM344). a** Kymographs from a representative cell showing L125-mNG fluorescence along the cell short axis (Y-axis) at midcell (top panel), L125-mNG fluorescence along the long axis (middle), and phase contrast signal along the cell long axis (bottom). **b** Representative phase contrast (left) and fluorescence (right) images of an L125-mNG expressing cell. In addition to the primary constriction at midcell, two secondary constrictions are visible at the cell quarter positions (white arrows). Scale bar, 1 µm. **c** Phase contrast intensity (top) and L125-mNG fluorescence (bottom) kymographs of the cell shown in panel b as a function of time. Time is relative to L125-mNG induction; dashed white lines indicate 0 min. Black arrows in the phase contrast kymograph indicate constrictions at the cell quarter positions, whereas those in the fluorescence kymograph indicate L125-mNG accumulation at the corresponding positions. **d** Length-binned kymograph of phase contrast intensity for the cell population after the start of L125-mNG induction (*N*=231). Black arrows indicate constrictions at the cell quarter positions.

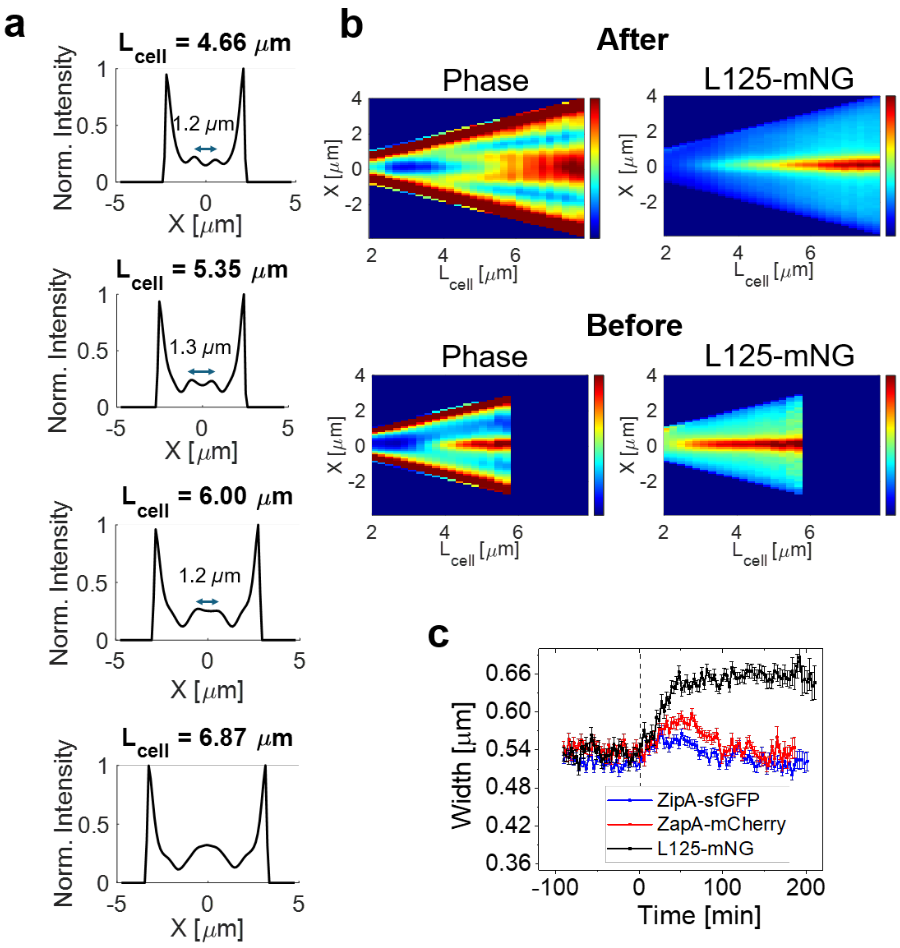


**Supplementary Fig. 5. The broad, shallow constrictions are much wider than the midcell accumulations of L125-mNG and the early Z-ring proteins. a** Average phase contrast intensity profiles along the long axis of cells at the indicated cell lengths, $L_{cell}$ after L125-mNG induction. The cell center corresponds to $X = 0$. The two local maxima near the cell center are separated by about 1.2-1.3 µm. **b** Population averaged cell length kymographs of phase contrast intensity (left) and L125-mNG (right) after L125-mNG induction (top) and before induction (bottom). Panels a,b use the same post-induction dataset (strain JM344, $N = 223$). The pre-induction dataset shown only in panel b comprises $N = 1070$. **c** Width of midcell accumulations of ZipA-sfGFP, ZapA-mCherry and L125-mNG fluorescence as a function of time. $N = 1336$ for ZipA-sfGFP (strain JM338), $N = 2494$ for ZapA-mCherry (strain JM333) and $N = 2127$ for L125-mNG (strain JM344) measurements. Error bars are s.e.m.

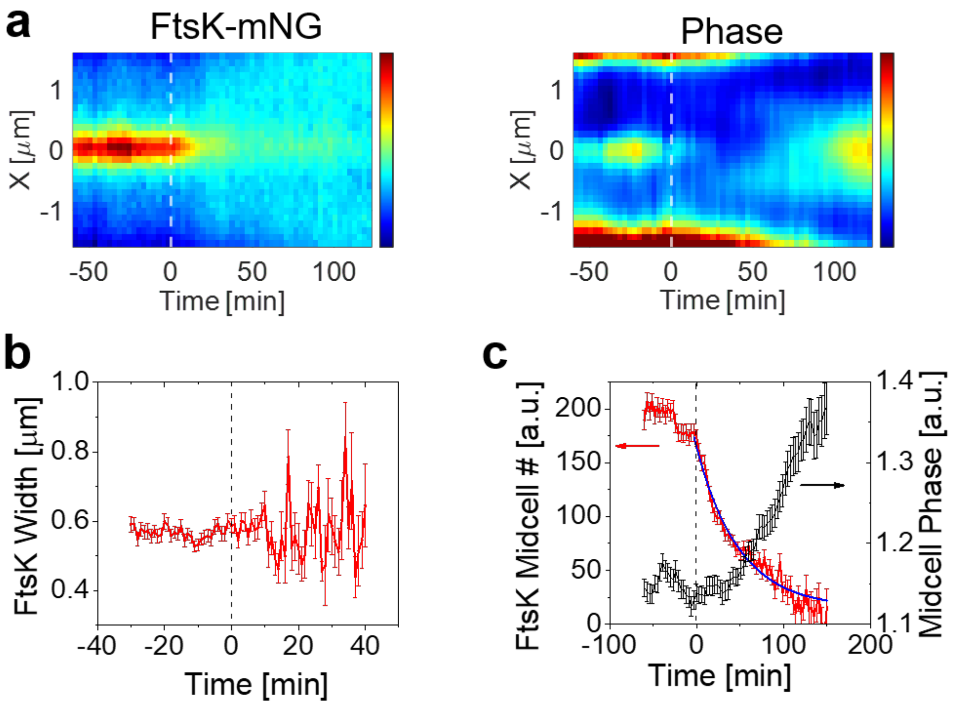


**Supplementary Fig. 6. FtsK localization after L125 induction. a** Population-averaged kymographs of FtsK-mNG signal (left) and phase-contrast intensity (right) before and after induction of L125 with 30 µM IPTG (strain JM356). Only the central region of the cells, extending ±1.5 µm from midcell, is shown. The dashed white line indicates the start of L125 induction. **b** Width of the midcell FtsK-mNG accumulation as a function of time relative to L125 induction. The width is determined by Gaussian fitting of FtsK-mNG intensity profiles along the cell middle (long axis). **c** Number of FtsK-mNG molecules at midcell (Z-ring) as a function of time relative to L125 induction (left axis). The numbers are averaged over the cell population and decay exponentially after L125 induction with a characteristic time of 49 min. Midcell phase-contrast intensity as a function of time for the same cells is shown on the right axis. *N*=845. Error bars in b and c are s.e.m.

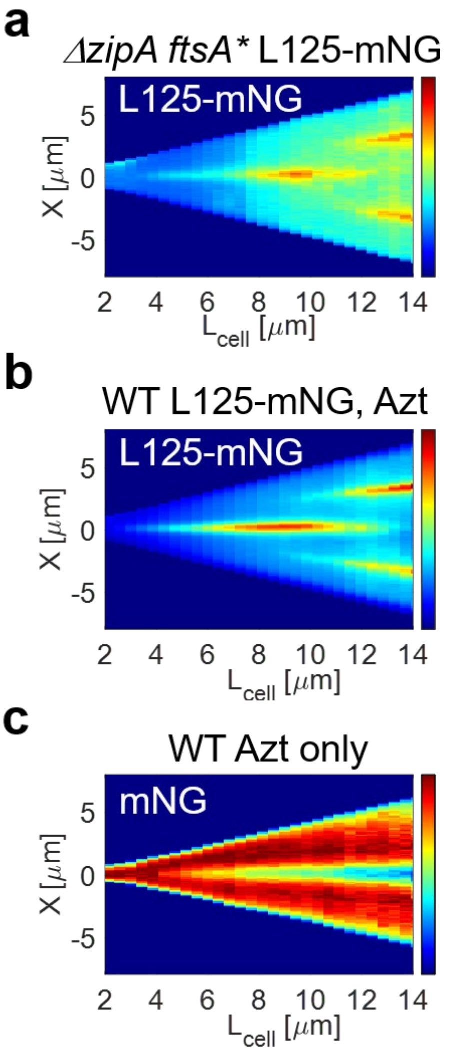


**Supplementary Fig. 7. ZipA is required for the formation of L125-induced constrictions, which persist during aztreonam treatment. a** Cell-length kymograph of L125-mNG fluorescence in *ΔzipA ftsA** cells (strain JM375) after induction of L125-mNG. *N*=312 cells. **b** Cell-length kymograph of L125-mNG fluorescence in wild-type cells (strain JM377) after simultaneous induction of L125-mNG and treatment with 0.3 µg/ml aztreonam (Azt). *N*=368 cells. **c** Cell-length kymograph of cytosolic mNG fluorescence in the control cells (strain JM228) treated with 0.3 µg/ml aztreonam. *N*=1022 cells. The cytosolic mNG signal serves as a marker of cell shape and the progression of constriction during aztreonam treatment.

**Supplementary Table 1. AlphaFold2 confidence metrics for *de novo* FtsA interface binder candidates.** Experimentally tested binders targeting the lateral or longitudinal FtsA interfaces are listed with binder length, pLDDT, pTM, interface pTM (i_pTM), normalized predicted aligned error (pAE), normalized interface predicted aligned error (i_pAE), interface pLDDT, and secondary-structure pLDDT (ss-pLDDT).

| Binder | Predicted FtsA interface | Length, aa | pLDDT | pTM | i_pTM | pAE | i_pAE | Interface pLDDT | ss-pLDDT |
|---|---|---|---|---|---|---|---|---|---|
| Lat62 | Lateral | 62 | 0.86 | 0.82 | 0.85 | 0.24 | 0.13 | 0.82 | 0.90 |
| Lat66 | Lateral | 66 | 0.91 | 0.84 | 0.85 | 0.22 | 0.12 | 0.89 | 0.93 |
| Lat123 | Lateral/ Longitudinal | 123 | 0.93 | 0.88 | 0.88 | 0.21 | 0.14 | 0.92 | 0.94 |
| L75 | Longitudinal | 75 | 0.89 | 0.91 | 0.87 | 0.23 | 0.26 | 0.86 | 0.90 |
| L81 | Longitudinal | 81 | 0.85 | 0.91 | 0.82 | 0.24 | 0.27 | 0.83 | 0.86 |
| L102 | Longitudinal | 102 | 0.93 | 0.92 | 0.91 | 0.17 | 0.21 | 0.92 | 0.94 |
| L125 | Longitudinal | 125 | 0.91 | 0.89 | 0.82 | 0.24 | 0.29 | 0.88 | 0.93 |

**Supplementary Table 2. Strains used in this study.**

| Strain | Relevant genetic marker(s) or features | Source, reference or construction |
|---|---|---|
| **BW27783** | Δ(araD-araB)567, ΔlacZ4787(::rrnB-3), Δ(araH-araF)570(::frt), ΔaraEp-532::frt, hsdR514, ϕPcp8araE535, rph-1, Δ(rhaD-rhaB)568, λ- | Yale Coli Genetic Stock Center (CGSC#: 12119) |
| **BW25113** | Δ(araD-araB)567, ΔlacZ4787(::rrnB-3), hsdR514, rph-1, Δ(rhaD-rhaB)568, λ- | Yale Coli Genetic Stock Center (CGSC#: 7636) |
| **W3110** | F-, λ-, IN(rrnD-rrnE)1, rph-1 | Lab stock/ Lutkenhaus lab |
| **DH5α** | F-, Φ80lacZΔM15, Δ(lacZYA-argF)U169, recA1, endA1, hsdR17(rk-, mk+), phoA, supE44, thi-1, gyrA96, relA1, λ- | NEB competent cells |
| **JM144** | BW27783<br>ΔftsN::frt-Ypet-ftsN<br>ΔzapA::zapA-mCherry-frt-cat-frt | (1) |
| **JM147** | BW27783<br>ΔftsZ::ftsZ$^{55-56}$-mNG<br>pEXT22 (empty vector) | (2) |
| **JM170** | BW25113<br>ΔftsI::frt-aph-frt-mNG-ftsI | λ Red recombineering |
| **JM171** | BW27783<br>ΔftsN::frt-mCherry-ftsN | Lab stock |
| **JM177** | BW27783<br>ΔzapA::zapA-mCherry-frt | (1) |
| **JM181** | BW27783<br>ΔftsN::frt-mCherry-ftsN<br>Δ(attL-lom)::bla lacIq P210-ftsA-GFP | P1 transduction of JM171 cells with P1vir lysate prepared from EC447 (3) |
| **JM182** | BW27783<br>ΔzapA::zapA-mCherry-frt-cat-frt<br>ΔftsI::frt-aph-frt-mNG-ftsI | P1 transduction of JM177 cells with P1vir lysate prepared from JM170 |
| **JM217** | BW25113<br>pSIM5 | (2) |
| **JM228** | BW27783<br>pEXT22-mNG | (2) |
| **JM269** | BW27783<br>ΔzipA::zipA$^{160-165}$-sfGFP<br>ΔyfeN::aph | P1 transduction of BW27783 cells with P1vir lysate prepared from CH128 (4) |
| **JM279** | BW27783<br>ΔzipA::zipA$^{160-165}$-sfGFP<br>ΔyfeN::frt | Kan$^R$ cassette excision using pCP20 |

| | | |
|---|---|---|
| **JM298** | BW27783<br>pJM-mNG (FtsA 5′ UTR) | Transformation of BW27783 |
| **JM301** | BW27783<br>pJM-Lat62 | Transformation of BW27783 |
| **JM302** | BW27783<br>pJM-Lat66 | Transformation of BW27783 |
| **JM303** | BW27783<br>pJM-Lat123 | Transformation of BW27783 |
| **JM304** | DH5α<br>pJM-Lat62-mNG | Transformation of DH5α |
| **JM305** | DH5α<br>pJM-Lat66-mNG | Transformation of DH5α |
| **JM306** | DH5α<br>pJM-Lat123-mNG | Transformation of DH5α |
| **JM320** | BW27783<br>pJM-L125 | Transformation of BW27783 |
| **JM333** | BW27783<br>ΔftsN::frt-Ypet-ftsN<br>ΔzapA::zapA-mCherry-frt-cat-frt<br>pJM-L125 | Transformation of JM144 |
| **JM334** | BW27783<br>ΔzapA::zapA-mCherry-frt-cat-frt<br>ΔftsI::frt-aph-frt-mNG-ftsI<br>pJM-L125 | Transformation of JM182 |
| **JM338** | BW27783<br>ΔzipA::zipA$^{160-165}$-sfGFP<br>ΔyfeN::frt<br>pJM-L125 | Transformation of JM279 |
| **JM342** | W3110<br>Δleu::Tn10, ftsA(R286W) (ftsA*)<br>pJM-L125 | Transformation of PS1321 (Lutkenhaus lab) |
| **JM343** | W3110<br>Δleu::Tn10, ftsA(R286W) (ftsA*)<br>ΔzipA::aph<br>pJM-L125 | Transformation of PS1322 (Lutkenhaus lab) |
| **JM344** | BW27783<br>pJM-L125-mNG | Transformation of BW27783 |
| **JM350** | DH5α<br>pJM-L75 | Transformation of DH5α |
| **JM351** | DH5α<br>pJM-L81 | Transformation of DH5α |

| | | |
|---|---|---|
| **JM352** | DH5α<br>pJM-L102 | Transformation of DH5α |
| **JM353** | BW25113<br>ΔftsX::ftsX-mNG-frt-aph-frt | λ Red recombineering |
| **JM354** | BW25113<br>ΔftsK::ftsK-mNG-frt-aph-frt | λ Red recombineering |
| **JM355** | BW25113<br>ΔftsX::ftsX-mNG-frt-aph-frt<br>pJM-L125 | Transformation of JM353 |
| **JM356** | BW25113<br>ΔftsK::ftsK-mNG-frt-aph-frt<br>pJM-L125 | Transformation of JM354 |
| **JM357** | TB28 (MG1655, lacIZYA::frt)<br>ΔftsB::aph<br>pBL194<br>pJM-L125 | Transformation of BL155/pBL194 (5) |
| **JM375** | W3110<br>Δleu::Tn10 ftsA(R286W) (ftsA*)<br>ΔzipA::aph<br>pJM-L125-mNG | Transformation of PS1322 (Lutkenhaus lab) |
| **JM377** | BW27783<br>pJM-L125-mNG-Kan$^R$ | Transformation of BW27783 |
| **JM385** | BW27783<br>pJM-L75-mNG | Transformation of BW27783 |
| **JM386** | BW27783<br>pJM-L81-mNG | Transformation of BW27783 |
| **JM387** | BW27783<br>pJM-L102-mNG | Transformation of BW27783 |

### Supplementary Table 3. Plasmids used in the study

| Plasmid | Relevant genetic marker(s) or features | Source, reference or construction |
|---|---|---|
| **pDSW210** | pBR-related ori; bla/Amp$^R$; lacIq; IPTG-inducible weak $P_{trc}$ | (3) |
| **pSEB306+** | pDSW210 derivative expressing full-length *ftsA*; bla/Amp$^R$ | (6) |
| **pJM21** | λ Red recombineering template for chromosomal C-terminal mNG fusions; mNG, FRT-aph-FRT, R6Kγ ori | Männik lab |
| **pJM139** | λ Red recombineering template for chromosomal N-terminal mNG fusions; mNG, FRT-aph-FRT, R6Kγ ori | Männik lab |
| **pEXT22-mNG** | IPTG-inducible plasmid for cytoplasmic mNG expression; aph/Kan$^R$ | (2) |
| **pEXT22** | IPTG-inducible plasmid aph/Kan$^R$ | (7) |
| **pJM-mNG** | pDSW210 derivative expressing mNG; bla/Amp$^R$ | This study |
| **pJM-Lat62** | pDSW210 derivative expressing codon-optimized binder Lat62; bla/Amp$^R$ | This study |
| **pJM-Lat62-mNG** | pDSW210 derivative expressing Lat62-mNG fusion; bla/Amp$^R$ | This study |
| **pJM-Lat66** | pDSW210 derivative expressing codon-optimized binder Lat66; bla/Amp$^R$ | This study |
| **pJM-Lat66-mNG** | pDSW210 derivative expressing Lat66-mNG fusion; bla/Amp$^R$ | This study |
| **pJM-Lat123** | pDSW210 derivative expressing codon-optimized binder Lat123; bla/Amp$^R$ | This study |
| **pJM-Lat123-mNG** | pDSW210 derivative expressing Lat123-mNG fusion; bla/Amp$^R$ | This study |
| **pJM-L75** | pDSW210 derivative expressing codon-optimized binder L75; bla/Amp$^R$ | This study |
| **pJM-L81** | pDSW210 derivative expressing codon-optimized binder L81; bla/Amp$^R$ | This study |

| | | |
|---|---|---|
| **pJM-L102** | pDSW210 derivative expressing codon-optimized binder L102; bla/$Amp^R$ | This study |
| **pJM-L75-mNG** | pDSW210 derivative expressing L75-mNG fusion; bla/$Amp^R$ | This study |
| **pJM-L81-mNG** | pDSW210 derivative expressing L81-mNG fusion; bla/$Amp^R$ | This study |
| **pJM-L102-mNG** | pDSW210 derivative expressing L102-mNG fusion; bla/$Amp^R$ | This study |
| **pJM-L125** | pDSW210 derivative expressing codon-optimized binder L125; bla/$Amp^R$ | This study |
| **pJM-L125-mNG** | pDSW210 derivative expressing L125 mNG fusion; bla/$Amp^R$ | This study |
| **pJM-L125-mNG-$Kan^R$** | pDSW210 derivative expressing L125 mNG fusion; aph/$Kan^R$ | This study |
| **pBL194** | pSC101ts, aadA/$Spec^R$, $P_{syn}$135::*gfp-ftsB* | (5) |
| **pSIM5** | pSC101-derived ori; repAts; cat/$Cam^R$; heat-inducible λ pL/CI857-*gam-exo-bet* | (8) |
| **pKD46** | oriR101; repA101$^{ts}$; bla/$Amp^R$; arabinose-inducible $P_{BAD}$-*gam-bet-exo* | (9) |
| **pCP20** | FLP recombinase expression; bla/$Amp^{R;}$ cat/$Cm^R$; temperature-sensitive replicon | (10) |

**Supplementary Table 4. Oligonucleotides used in this study.**

| Name | Sequence (5'->3') | Purpose |
|---|---|---|
| PR219 | TGGACGAACTGTACAAGTAAAAGCTTCTGTTTTGGCGG | Universal oligonucleotide for generating C-terminal mNG fusions to binder constructs |
| PR220 | TCACCTTTGCTGACGAATTCCTGCGAATGCATATAGATGTC | C-terminal mNG fusion to Lat62 |
| PR221 | TCACCTTTGCTGACGAATTCGTGATGACGAATCAACTCATC | C-terminal mNG fusion to Lat66 |
| PR222 | TCACCTTTGCTGACGAATTCGCTGCGTTTAGCACGGTTTG | C-terminal mNG fusion to Lat123 |
| PR223 | TCACCTTTGCTGACGAATTCTTTCATATACTTAGGAATCAAT GCAATC | C-terminal mNG fusion to L125 |
| PR224 | TCACCTTTGCTGACGAATTCTCCAATGGTGCTCAGCCATTTT TC | C-terminal mNG fusion to L75 |
| PR225 | TCACCTTTGCTGACGAATTCCCACTTTTCCAGATAAGGTG | C-terminal mNG fusion to L81 |
| PR226 | TCACCTTTGCTGACGAATTCTTTCTTGTATTTTTCAATTGCC TTC | C-terminal mNG fusion to L102 |
| PR227 | GAATTCGTCAGCAAAGGTG | Amplification of mNG for HiFi assembly |
| PR228 | TTACTTGTACAGTTCGTCC | Amplification of mNG for HiFi assembly |
| PR229 | AGAAAATATCGTAGTGCAAAAATAAGGATAAACGCGACG CTGTAGGCTGGAGCTGCTTCG | λ Red recombineering of chromosomal *ftsI* for mNG-FtsI |
| PR230 | CTTCCTGACGTTTTGGTTTCTGCGTTTTCGCCGCTGCTTTCG CGCTGCCAGAACCAGCGG | λ Red recombineering of chromosomal *ftsI* for mNG-FtsI |
| PR231 | GCTTGCCACGGTACAACATTTACGCCACTTTACGCCTGAAA GCTCGGCTGGCTCCGCTG | λ Red recombineering of chromosomal *ftsX* for FtsX-mNG |
| PR232 | TACGGAACCCATTGCAGGGAAAGAGTATAACACGCTTTTA CATATGAATATCCTCCTTAG | λ Red recombineering of chromosomal *ftsX* for FtsX-mNG |
| PR233 | CAACGGTAATCGTGAAGTGCTGGCCCCACCGCCGTTTGAC AGCTCGGCTGGCTCCGCTG | λ Red recombineering of chromosomal *ftsK* for FtsK-mNG |
| PR234 | TGCGACGCTACCGCGCCTTATCCGGCATACGATGCATTAG CATATGAATATCCTCCTTAG | λ Red recombineering of chromosomal *ftsK* for FtsK-mNG |
| PR235 | TTTACGCCACTTTACGCCTG | Verification of chromosomal FtsX-mNG fusion |
| PR236 | TCCACAAGTTCAATGCAAGCT | Verification of chromosomal FtsX-mNG fusion |
| PR237 | CGGTAATCGTGAAGTGCTGG | Verification of chromosomal FtsK-mNG fusion |

| | | |
|---|---|---|
| PR238 | TCAGTTGATAGAGTGCCGGA | Verification of chromosomal FtsK-mNG fusion |
| PR239 | GTTCTTCTGACTGTCAGACCAAGTTTAC | Amplification of pJM-L125-mNG lacking the *amp*$^R$ cassette for HiFi assembly |
| PR240 | GTTCAATCATACTCTTCCTTTTTCAATATTATTG | Amplification of pJM-L125-mNG lacking the *amp*$^R$ cassette for HiFi assembly |
| PR241 | AAGGAAGAGTATGATTGAACAAGATGGATTG | Amplification of the *kan*$^R$ cassette for HiFi assembly |
| PR242 | GGTCTGACAGTCAGAAGAACTCGTCAAG | Amplification of the *kan*$^R$ cassette for HiFi assembly |

**Supplementary Table 5. Coding sequences of experimentally tested FtsA interface binders.**

Binder-coding sequences were codon-optimized for expression in *E. coli* using the IDT Codon Optimization Tool, and an ATG start codon was introduced where required.

### Lat62

**Amino-acid sequence:**
MVDKKEAYQKQYRLWEEYQASEEKGRSFWWYMDKQVRRGNPDARVVVRYYKEYHPDIYMHSQ

**Codon-optimized coding sequence:**
ATGGTCGACAAAAAGGAAGCGTATCAAAAGCAGTATCGCTTGTGGGAGGAATATCAGGCTTCGGAAGAAAAGGGTCGTTCAT
TCTGGTGGTATATGGACAAACAAGTACGCCGTGGTAACCCCGATGCTCGCGTAGTGGTACGTTACTACAAAGAATACCACCCG
GACATCTATATGCATTCGCAGTAA

---

### Lat66

**Amino-acid sequence:**
MEKIRVLREYIPTLTEITKNSKSSSIYSQSRRLVEEMEEFIRQGKPITEKRFRSYKYGADELIRHH

**Codon-optimized coding sequence:**
ATGGAAAAGATTCGTGTGTTACGTGAGTATATCCCGACCCTGACAGAGATTACTAAAAACAGCAAGTCAAGCTCGATCTACAG
CCAGAGCCGCCGCCTTGTGGAAGAGATGGAGGAATTTATTCGTCAAGGAAAGCCCATTACCGAAAAACGCTTTCGCTCGTATA
AGTACGGTGCGGATGAGTTGATTCGTCATCACTAA

---

### Lat123

**Amino-acid sequence:**
MKEERVIRMIDSGDNSPSTVNTISMEVSTIMIHHAMKVDLSPSVKKRMRKADATWIAAYNESKRTGFTHEIYRTMYEATEEMRVV
MEEHVYSKMSPEQAEIQKYLVTHVMAWLKEEANRAKRS

**Codon-optimized coding sequence:**
ATGAAGGAAGAACGTGTAATTCGTATGATTGACAGCGGTGATAATAGCCCGAGCACAGTGAACACCATTAGCATGGAAGTAT
CTACAATCATGATCCATCACGCGATGAAGGTCGACTTAAGCCCTAGCGTAAAAAAACGCATGCGTAAAGCAGATGCGACGTGG
ATTGCAGCGTATAACGAAAGCAAACGTACCGGCTTCACTCACGAGATTTACCGCACCATGTATGAGGCGACAGAAGAAATGCG
CGTGGTGATGGAAGAGCACGTATATTCTAAAATGTCGCCAGAACAAGCAGAAATTCAGAAATACCTGGTAACACACGTTATGG
CCTGGCTGAAGGAAGAAGCAAACCGTGCTAAACGCAGCTAA

---

## L75

**Amino-acid sequence:**
MEEFRHKAAEVFHKFNTAWFYHFFRGDEEKTEELDKEWERLMPLMIEARKTFEKGDLTPLEEAKAIAEKWLSTIG

**Codon-optimized coding sequence:**
ATGGAAGAATTTCGCCATAAGGCAGCGGAAGTGTTTCATAAGTTCAATACCGCGTGGTTTTACCATTTCTTTCGCGGCGACGAA
GAAAAAACGGAAGAACTTGATAAGGAATGGGAACGTCTGATGCCTCTGATGATTGAGGCCCGTAAAACCTTCGAGAAGGGCG
ATCTGACCCCGCTGGAAGAAGCCAAAGCCATCGCGGAAAAATGGCTGAGCACCATTGGATAA

---

## L81

**Amino-acid sequence:**
MSERELIIQVFYLHDVMIMSGYWYDPELQELYDEVFEIMLELKKNKDNPEKRKELIEKIKEKFEKLKELLEKHPSPYLEKW

**Codon-optimized coding sequence:**
ATGAGCGAACGTGAATTAATCATTCAGGTCTTTTACTTACACGATGTTATGATTATGTCTGGTTACTGGTACGACCCGGAATTG

CAGGAACTGTATGACGAGGTATTTGAAATCATGCTGGAACTGAAAAAAAACAAAGATAATCCAGAAAAACGCAAGGAACTGA
TCGAAAAGATCAAAGAAAAGTTCGAGAAGTTAAAGGAACTGTTGGAGAAACATCCGTCACCTTATCTGGAAAAGTGGTAA

---

## L102

**Amino-acid sequence:**
MPLIEKLEELVEGLSALWERIKNREITDPAEIKAQLEPLFEKALPVLEELLKEKVMSEREVELLREEIEYVSSDEVFPRWYLRDQAFFFAH
VFKKAIEKYKK

**Codon-optimized coding sequence:**
ATGCCCCTGATTGAAAAATTAGAAGAATTAGTGGAGGGGCTGTCGGCCCTGTGGGAACGCATTAAAAATCGCGAGATTACGG
ACCCGGCGGAAATTAAAGCGCAGTTGGAACCGCTGTTTGAAAAAGCCCTGCCCGTTTTGGAAGAATTATTGAAGGAAAAAGT
GATGAGCGAGCGTGAAGTAGAACTGCTGCGTGAGGAGATTGAGTATGTCTCGTCTGATGAAGTATTTCCGCGCTGGTATCTGC
GTGATCAAGCCTTTTTTTTCGCGCATGTTTTCAAGAAGGCAATTGAAAAATACAAGAAATAA

---

## L125

**Amino-acid sequence:**
SAALEEILRRAEALRREVEELLGLLEEGVEISLEELRERLEALTARAREVDHLVWANFKDSHPELYEEATHIYWSLDHVLTMFETFLTKP
GRPSDEFIRKWVALNLRYVLAEVDELIALIPKYMK

**Codon-optimized coding sequence:**
ATGAGTGCCGCTCTGGAAGAAATCCTGCGTCGTGCTGAAGCGCTGCGCCGCGAAGTTGAAGAATTGTTGGGCTTACTGGAGG
AGGGTGTGGAAATTTCCCTTGAGGAACTGCGTGAACGCCTGGAAGCACTTACAGCCCGCGCACGCGAGGTCGACCATCTGGT
TTGGGCCAACTTTAAAGATAGCCATCCAGAACTGTATGAAGAAGCGACGCATATCTACTGGTCTCTGGATCACGTACTGACAAT
GTTTGAAACGTTTCTGACCAAACCGGGTCGCCCGAGCGATGAATTTATTCGCAAGTGGGTGGCACTGAATCTTCGTTATGTCCT
GGCGGAGGTGGATGAGCTGATTGCATTGATTCCTAAGTATATGAAATAA

---
